# Within-host evolution of SARS-CoV-2: how often are *de novo* mutations transmitted?

**DOI:** 10.1101/2023.08.08.552503

**Authors:** Chapin S. Korosec, Lindi M. Wahl, Jane M. Heffernan

## Abstract

Despite a relatively low mutation rate, the large number of SARS-CoV-2 infections has allowed for substantial genetic change, leading to a multitude of emerging variants. Using a recently determined mutation rate (per site replication), as well as parameter estimates for within-host SARS-CoV-2 infection, we apply a stochastic transmission-bottleneck model to describe the survival probability of *de novo* SARS-CoV-2 mutations. For narrow bottlenecks, we find mutations affecting pertarget-cell attachment rate (with phenotypes associated with fusogenicity and ACE2 binding), have similar transmission probabilities to mutations affecting viral load clearance (with phenotypes associated with humoral evasion). We further find that mutations affecting the eclipse rate (with phenotypes associated with reorganization of cellular metabolic processes and synthesis of viral budding precursor material) are highly favoured relative to all other traits examined. We find mutations leading to reduced removal rates of infected cells (with phenotypes associated with innate immune evasion) have limited transmission advantage relative to mutations leading to humoral evasion. Predicted transmission probabilities, however, for mutations affecting innate immune evasion are more consistent with the range of clinically-estimated household transmission probabilities for *de novo* mutations. This result suggests that although mutations affecting humoral evasion are more easily transmitted when they occur, mutations affecting innate immune evasion may occur more readily. We examine our predictions in the context of a number of previously characterized mutations in circulating strains of SARS-CoV-2. Our work offers both a null model for SARS-CoV-2 substitution rates and predicts which aspects of viral life history are most likely to successfully evolve, despite low mutation rates and repeated transmission bottlenecks.

## 1 Introduction

The emergence and rapid spread of severe acute respiratory syndrome coronavirus 2 (SARS-CoV-2) in late 2019 marked the onset of an unprecedented global health crisis. The evolution of SARS-CoV-2 variants of concern (VOC) has since posed immense challenges to public health, impacting societies world-wide. A mutant lineage nomenclature based on phylogenetic framework was developed, which enabled tracking single nucleotide polymorphisms (SNPs), as well as linking specific SNPs to changes in viral fitness traits [1]. Currently, the XBB.1.5 variant, a recombinant of two BA.2 Omicron sub-variants, is the predominant variant of concern circulating globally. XBB.1.5 has ‘high’ growth advantage and ‘moderate’ ability for immune escape as compared to previous omicron subvariants, as determined by The World Health Organization (WHO) [2].

The mutation rate of SARS-CoV-2 has been estimated using genome sequencing and phylogenetics, and is relatively low for viral pathogens [3]. Since case-counts worldwide exceed 750 million cumulative infections to date [4], however, SARS-CoV-2 has accrued substantial genetic diversity [5]. At the population level much attention has been focused on quantifying between-host diversity and determining current SARS-CoV-2 epidemiological traits [6–8]. However, novel mutations first occur within a single individual, and thus methodology capable of predicting within-host diversity, and forecasting the transmission probability for novel mutations is essential for predicting the emergence of future SARS-CoV-2 VOC. The ongoing threat to public health from SARS-CoV-2 evolution necessitates a need for “a full model incorporating within-host evolutionary dynamics and transmission” [9] to predict how new mutations arise and are transmitted, and to predict the rate at which new variants will arise, so that best-informed interventions can be enacted.

We have developed a data-driven model describing SARS-CoV-2 within-host evolutionary dynamics. The model is used to compute the probability that at least one copy of a novel mutation arising in a host survives to be transmitted to a new host. We quantify the fate of neutral mutations, as well as mutations leading to immune escape, increased transmissibility, binding affinity, and replication. For each mutant lineage, we explore how bottleneck size and mutant fitness affect the transmission probability. Finally, we contextualize our model predictions with data from known SARS-CoV-2 mutations that have been quantitatively linked to phenotypic changes in viral fitness.

## 2 Methods

Within-host modelling of SARS-CoV-2 has helped to elucidate the molecular mechanisms driving SARS-CoV-2 transmission as well as immunological responses from vaccines and infection [10–16], as well as offer key insights into the potential implications of public health burden and interventions [17–19]. In this work, our within-host model is similar in structure to models of within-host infection that have been used to quantify the dynamics of SARS-CoV-2 within an infected individual [20–22], and estimate viral fitness traits for a number of SARS-CoV-2 VOCs (i.e., attachment rate, budding rate) [20, 23–26]

A schematic of our in-host model of SARS-CoV-2 infection with three eclipse stages (Eq S1) is shown in Fig 1A. A schematic of the stochastic model employed to determine transmission probabilities of rare mutations is shown in Fig 1B. Parameters for “wildtype” SARS-CoV-2 infection (see Section S1.2 and Discussion) are determined from the literature and are listed in Table S1.

**Fig. 1.**
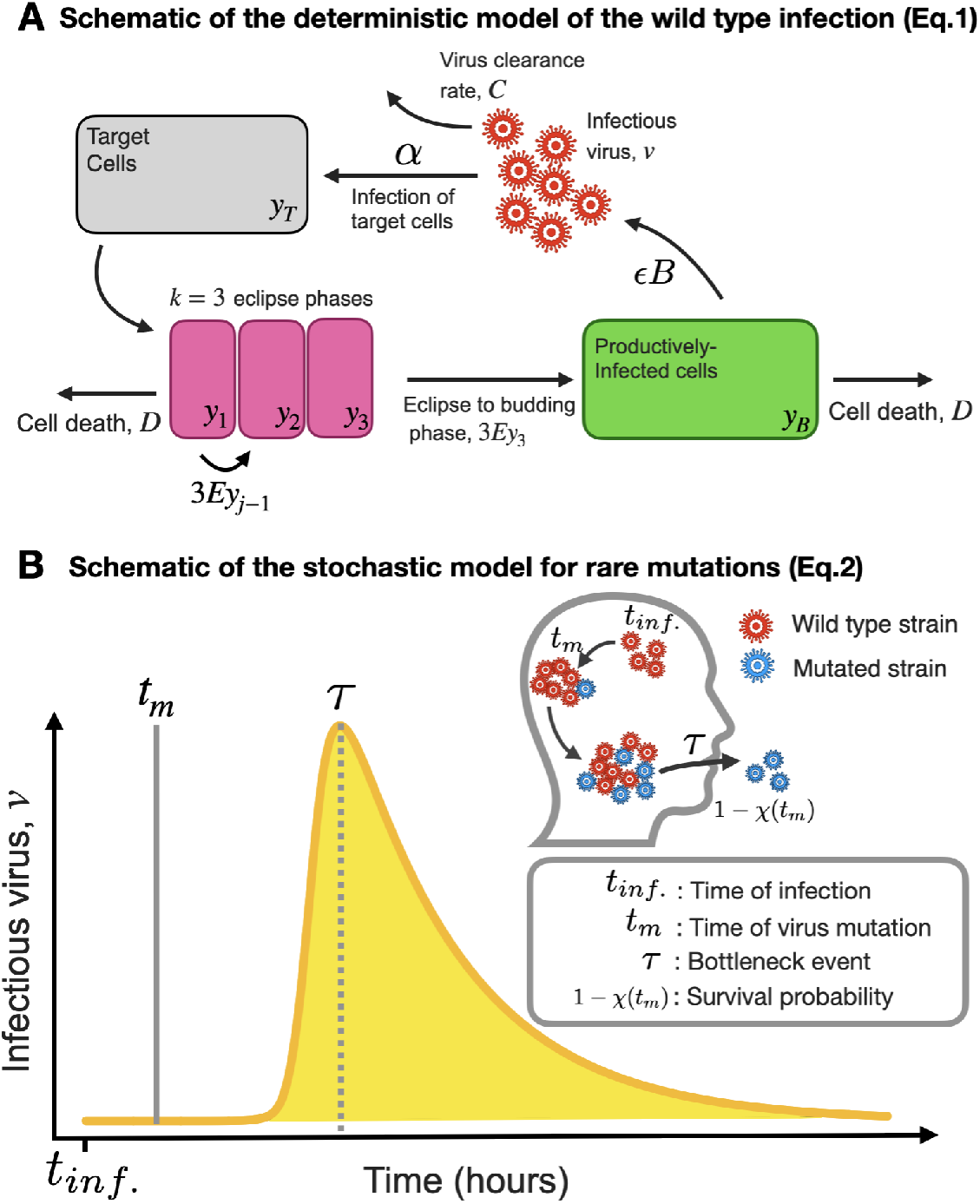
A) Schematic of the deterministic model of the wildtype infection (Eq. S1) described in Section S1.1. B) Schematic of the stochastic model for rare mutations (Eq. S2) described in Sections S1.3 and S1.6.

The within-host diversity of SARS-CoV-2 is relatively low, with narrow transmission bottlenecks [9, 27–29]. This small bottleneck size leads to low diversity at the time of transmission, whereby the majority variant is typically transmitted to a new host [29], and thus the emergence and spread of a *de novo* mutation will be a statistically rare event. While modelling and prediction of rare, impactful events can be approached through computational simulations [30], we follow ref.[31] in developing a rigorous within-host mathematical model of the underlying process. This allows for a full analytical exploration of the factors driving the emergence of a VOC. Following [31], we develop a model for the wildtype population dynamics (Section S1.1) and a stochastic model for the introduction of new and rare viral variants (Section S1.3). Model parameter values are informed from the literature (Section S1.2). To capture the emergence of a new mutant and the growth of the mutant and wildtype population, we couple the stochastic and deterministic models (Section S1.4). Using the percent change in the Basic Reproduction Number as a metric of fitness of de novo advantageous mutants, we define life-history and immune escape traits that can affect the emergence and transmissibility of a mutant strain (Section S1.5). Finally, we determine the probability that a new variant that arises at time 0 *< t*_*m*_ *< τ* can be transmitted at time *τ*, the time of the bottleneck event (Section S1.6) which is assumed to occur at peak viral load.

We note that the modelling framework presented here does not explicitly model the immune response. The immune response is instead included in model parameters, for example, reflecting virus neutralization by antibodies (*C*), infected cell killing by cytotoxic T-lymphocytes (*D*), and the ability of the virus to infect a target cell modified by interferon (*α*). Furthermore, we make the simplifying assumption that these parameters do not vary with time. This is reasonable since we model the in-host infection only to the point of peak viral load, and the adaptive immune response typically plays a limited role before this time [32].

## 3 Results

### 3.1 Neutral strain bottleneck dynamics for narrow bottlenecks and selective coefficient

A schematic of the deterministic model with three eclipse stages (Eq S1) is shown in Fig 1A, and a schematic of the stochastic model is shown in Fig 1B. The SARS-CoV-2 widltype strain parameters are determined from the literature and are listed in Table S1; the infection time-course for the wildtype strain is shown in Fig. 2A.

**Fig. 2.**
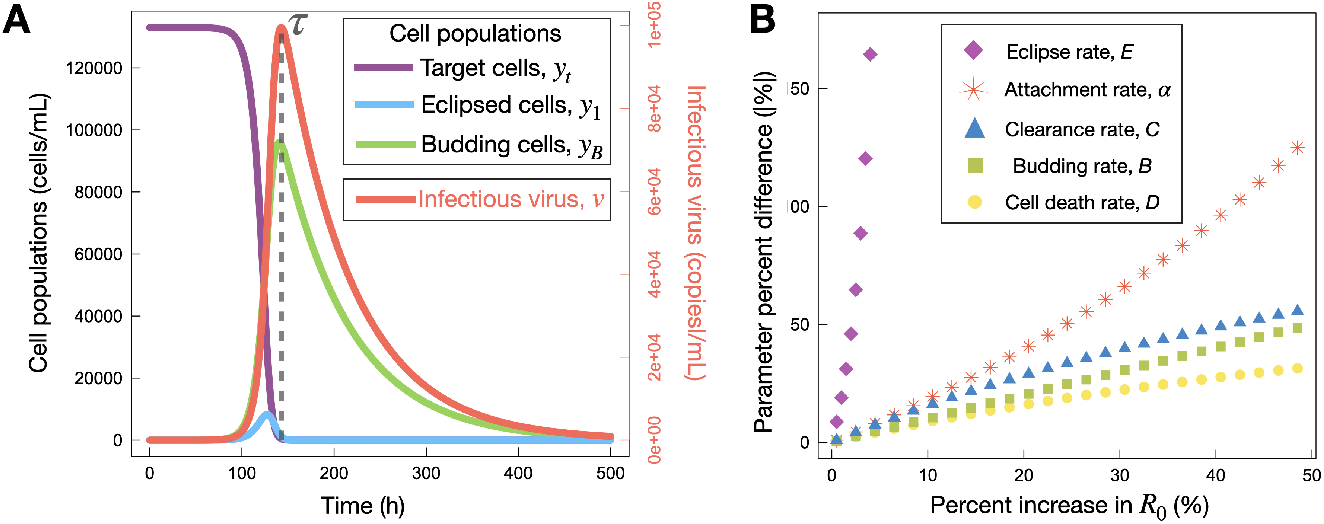
Wildtype dynamics. A) Timecourse of the wildtype infections (Eq. S1). Cell populations are plotted against the left-hand axis, while infectious virions are plotted against the right-hand axis. The grey dashed line indicates the transmission bottleneck time, *τ*, which is fixed for all transmission events to the time of peak infectious virus. B) Percent change in model parameters required to achieve a specific percent increase in the reproductive ratio, *R*_0_. Wildtype parameters are listed in Table S1.

The infection time-course for the wildtype strain is shown in Fig. 2A. The rare mutations studied in the stochastic model can affect any of five viral traits: attachment, budding, viral clearance, infected cell death, and eclipse phase timing, captured in parameters *α, B, C, D* and *E* respectively. Fig. 2B illustrates the percent change in the wildtype parameter value that is required to achieve a specific increase in *R*_0_ for the variant strain. We note that very large increases in the eclipse rate *E* (reflecting substantial reductions in the eclipse phase duration) are needed to achieve even moderate changes in *R*_0_.

Fig.S1A and Fig.S1B display the survival probability of a *de novo* mutation (*P*_*s*_, Eq. S5) and the rate at which surviving lineages are created (*κ*, Eq. S6) respectively, for neutral mutations. We note that the neutral strain reproduces the ‘hockey stick’ pattern found for neutral mutations in Influenza A [31], as shown in the semi-log plot of *P*_*s*_ (Fig.S1B).

### 3.2 Variant dynamics with narrow bottlenecks (Λ = 1)

We begin by considering the fate of rare mutations for narrow bottlenecks of size Λ = 1. As seen in Fig 3 (*y*-intercept), we find that the probability of transmitting at least one mutant (Π, Eq. S8), is approximately 5.2×10^−7^ for a neutral mutation with a transmission bottleneck of size one.

**Fig. 3.**
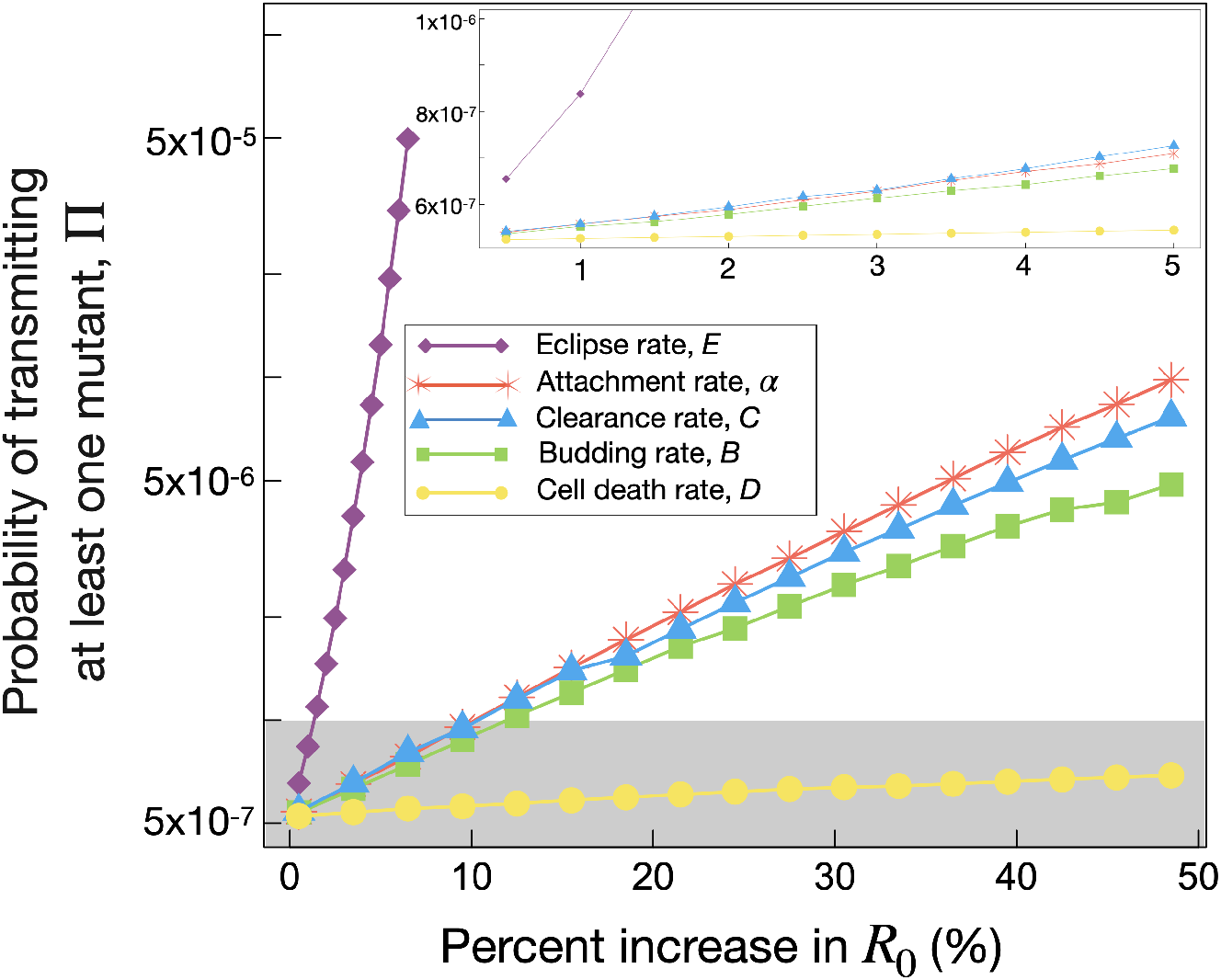
Probability of transmitting at least one mutant, Π (Eq. S8), as a function of percent increase in *R*_0_ (Eq. S4), for a bottleneck of size Λ = 1. Results for mutations affecting parameters *E, α, C, B*, and *D* are shown in purple, red, blue, green, and yellow, respectively. Grey box and inset show experimental parameter ranges in Π (see Discussion).

Coloured lines in Fig 3 predict Π for variants that carry adaptive (beneficial) mutations affecting different traits. We see that mutations affecting the eclipse phase timing have a high transmission probability, relative to other traits, while mutations affecting the death rate of infected cells are least likely to survive transmission. Mutations in *α, C*, and *B* lead to a similar increase in Π as a function of *R*_0_ We note that mutations in *E* are limited to a maximum increase in 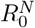 of 6.5%, as further increases in 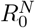 would require extreme changes in eclipse phase timing (see Fig. 2B).

For completeness, Fig.S1C and Fig.S1D display *P*_*s*_ (Eq. S5) and *κ* (Eq. S6) for mutations leading to a 5% 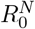 increase, for each model parameter. All mutations lead to an increase in *P*_*s*_ and *κ* for all *t*_*m*_ as compared to neutral mutations. Fig.S2 and Fig.S3 show the increase in *P*_*s*_ and *κ*, respectively, for all mutation parameters and all selective coefficients.

### 3.3 Changes in variant dynamics with transmission bottleneck size

As described in the Introduction, estimates of the transmission bottleneck size for SARS-CoV-2 include a range of values. Fig. 4 displays Π as a function of Λ for mutations leading to an increase in *R*_0_ of 0.5% (Fig. 4A), 5.0% (Fig. 4B), or 50% (Fig. 4C). Overall, we see that the probability that a new variant is transmitted increases with bottleneck size.

**Fig. 4.**
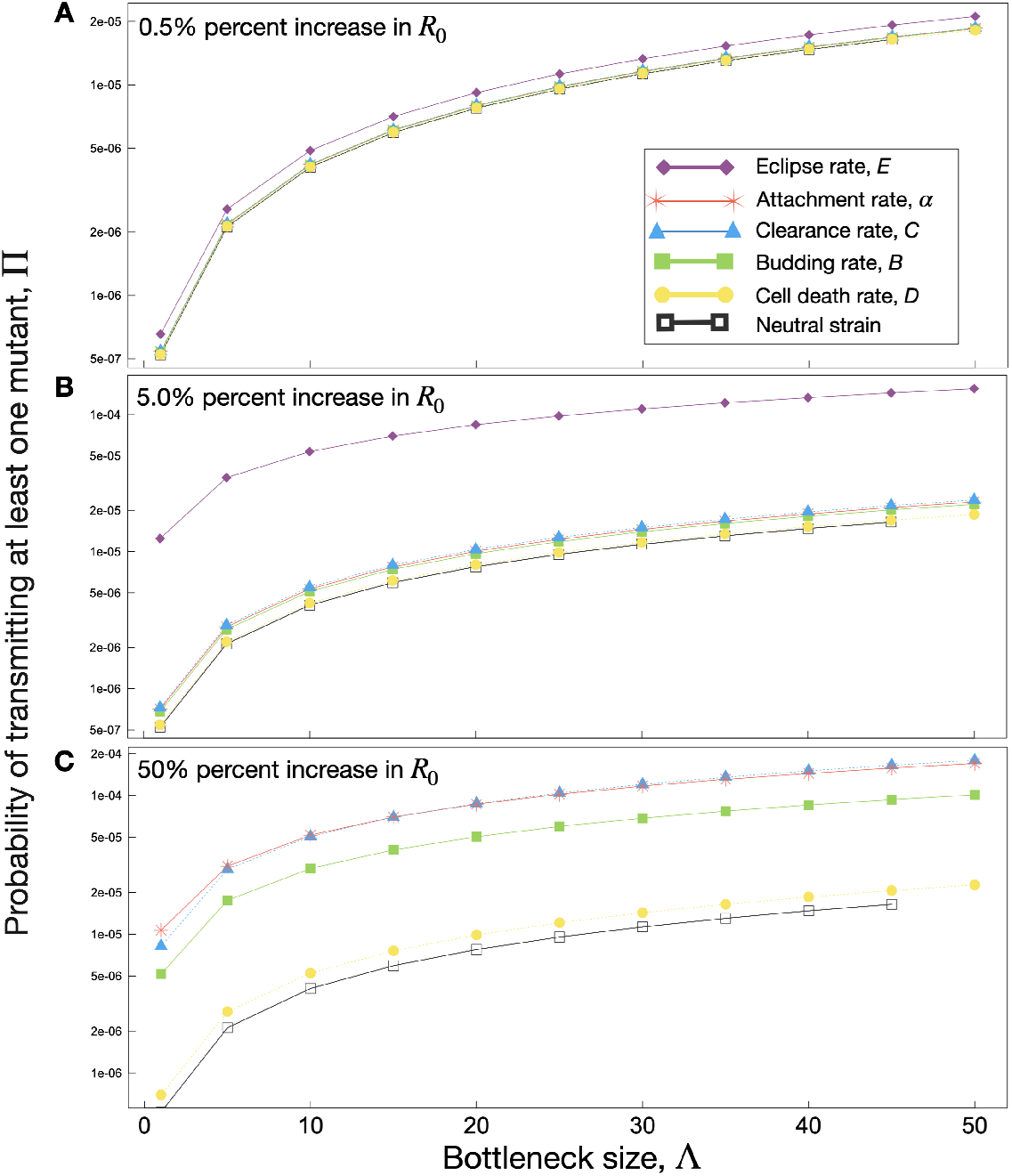
Probability of transmitting at least one mutant, Π (Eq. S8), as a function of bottleneck size for selective coefficients 0.5% (panel A), 5% (panel B), and 50% (panel C). *E* is omitted from panel C as only selective coefficients up to 6.5% were examined for this parameter.

An *R*_0_ increase of 0.5% leads to a similar increase in Π with Λ for all mutation types, with transmission probabilities close to those expected for a neutral mutation. When the variant has a 5.0% increase in *R*_0_, a clear hierarchy of parameter influence on Π as a function of Λ emerges. As seen before, mutations affecting the eclipse phase timing are the most likely to be transmitted, irrespective of the bottleneck size. For very fit variants (50% increase in *R*_0_), eclipse phase mutations are no longer realistic as described previously. Here, mutations affecting the attachment rate *α* or clearance rate *C* are found to have a higher probability of transmitting. Mutations in the budding rate *B* have an intermediate effect on Π, while mutations affecting the cell death rate, *D*, appear to offer a limited transmission advantage compared to the neutral strain. This final result is surprising, given that the mutation in question confers a 1.5 times increase in *R*_0_ when compared to the wildtype. To further illustrate the selective advantage of each mutation over the neutral strain, in Fig.S4 we report the ratio of the mutant Π to that of the neutral strain for each parameter. Notably, we find that with a selective coefficient of only 5.0%, *E* has a 20x higher probability of transmitting at narrow bottlenecks, whereas the selective coefficient needs to be as high as 50% for mutations in *α* and *B* to have a 20x higher probability of transmitting at narrow bottlenecks. In Fig. S5 we examine these trends in greater detail, combing results for Π as a function of both *R*_0_ and bottleneck size. We see that the increase in Π is roughly exponential with mutant fitness, for mutations affecting *α, B, C* and *D*, and find that Π scales faster than exponentially as a function of *R*_0_ for mutations affecting *E*.

## 4 Discussion

Pathogens face significant selective pressure to proliferate within the host, whereas host defensive mechanisms are geared towards impeding pathogen replication. This ongoing conflict between the host and pathogen creates an arms race, resulting in diverse outcomes that can either enhance or diminish virulence [33]. In this study, we used a stochastic model to explore the fate of neutral and advantageous SARS-CoV-2 mutations, where the wildtype strain was calibrated to SARS-CoV-2 within-host viral load data collected from February 2020 through to February 2021 in Berlin, Germany [21], with model parameters determined through previous work [23].

Our computed probabilities of transmitting at least one mutant, Π, compare well to recent estimates of SARS-CoV-2 transmission probabilities from both clinical and simulation studies. In a deep sequencing study of clinical isolates, Lythgoe *et al*. [9] report that the mean number of intrahost single nucleotide variants (iSNVs) observed per 100 sites is 4×10^−3^, implying that the probability of an iSNV becoming established, in a single patient at a particular site, is 4×10^−5^. They further report that approximately 400 iSNVs were observed in households, of which four appeared to be transmitted (Figure 3B of ref. [9]). Thus, a rough estimate is that 1/100 iSNVs are transmitted after having been established in the donor. Based on these estimates, the probability that a new mutation occurs at a particular site and is then transmitted is approximately 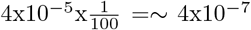, in close agreement with the analytical estimates we report here. In other work, Van Egeren *et al*. [30] explore the probability of variant transmission as a function of increased infection period through intrahost viral dynamic simulations. They find that approximately 3-4 weeks post initial infection, the probability of transmitting a single mutated virion is on the order of 10^−7^*−* 10^−6^ (Fig 3C from ref. [30]). Thus, both clinical and simulation results compare remarkably well to our data-driven analytical estimates at narrow bottlenecks (Λ*≤* 5).

If we restrict our results to narrow bottlenecks (Λ = 1), with transmission probabilities confined to the range 10^−7^ *−* 10^−6^, the grey region and inset of Fig. 3 depicts the predicted parameter space for mutants affecting each of the examined parameters and also illustrates the maximum increases in selective coefficients of key viral life-history traits. We can see that all mutations reducing infected cell death, *D*, are within this region. In contrast, mutations affecting the eclipse rate are only included for a maximum increase in *R*_0_ of *∼*2.5%. Mutations affecting *α, C*, and *B* display a similar response as a function of percent increase in *R*_0_, and are intermediate to *D* and *E*. Using 1*×*10^−6^ as a maximum transmission probability, we can also place limits on the percent change relative to the neutral strain of each of our parameters, leading to a prediction for limitations to likely phenotypic changes caused by a single advantageous mutant. Relative to the neutral strain, we find that parameters *α, B, C, D*, and *E* maximally shift *R*_0_ by 18.4, 18.9, 11.5, 32.0, and 15.5%, respectively. Thus, by bounding the probability of transmission to 1 *×*10^−6^ as suggested by clinical data [9], we can put an upper bound on changes in key phenotypic traits that have arisen from a single mutation.

Next, we focus on mutations affecting each of these traits individually, examining our predictions in the context of identified SARS-CoV-2 mutations. The predominant structural proteins of SARS-CoV-2 include membrane glycoprotein (M), envelope protein (E), nucleocapsid protein (N), and the spike protein (S) [34]. Mutations in M, E, N, or S can affect various life-history traits captured by our stochastic model describing the fate of mutant lineages (Eq. S2). We briefly summarize a number of key mutations identified in SARS-CoV-2 VOC in Table 1, with reference to expected changes in our model parameters based on observed phenotypic effects.

**Table 1.**
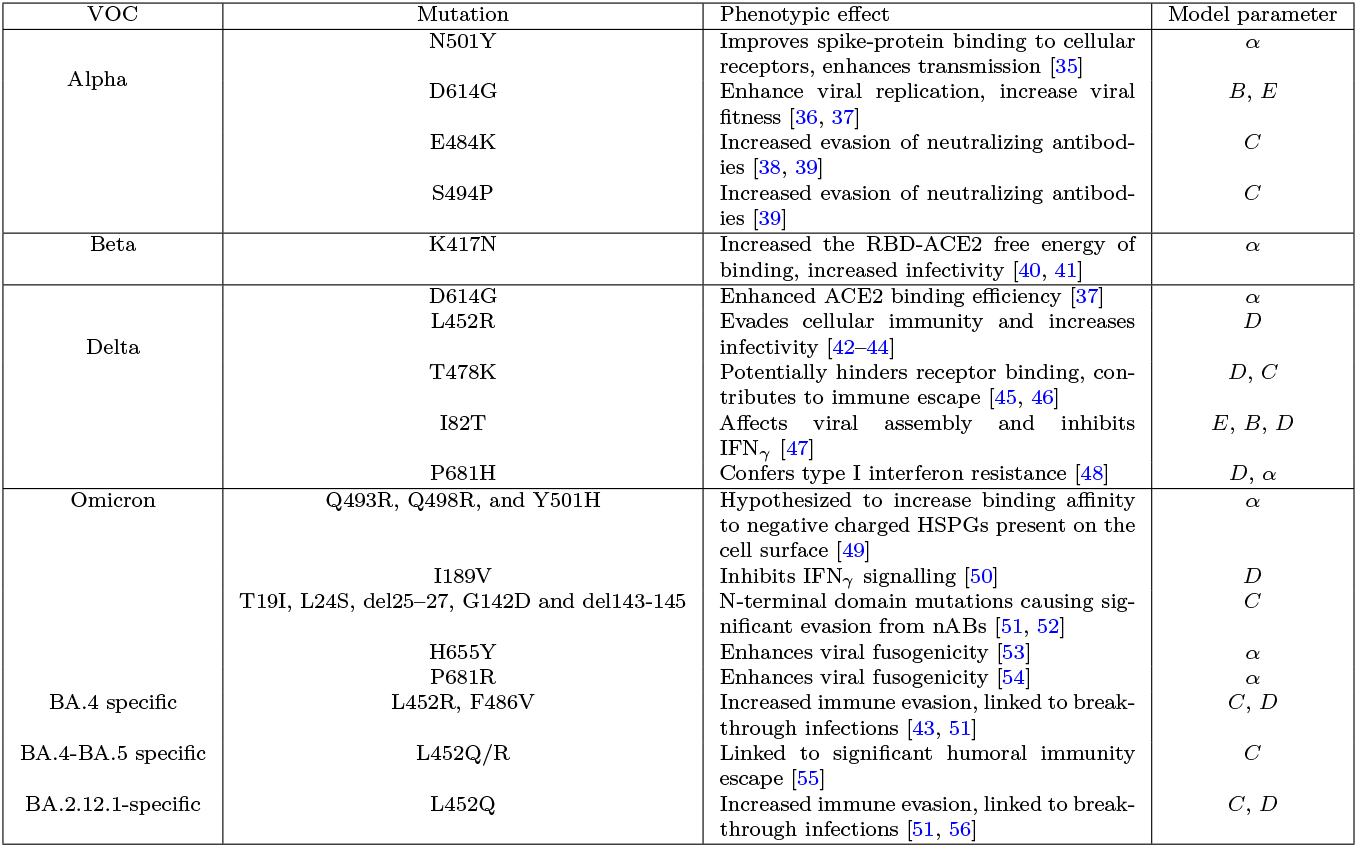
Summary of mutations, phenotypic effects, and associated predicted change to within-host model parameter(s).

The per target-cell attachment rate, *α*, represents a myriad of interactions involved in a virus gaining access to the host-cell intracellular environment [57]; from initial binding, diffusion throughout the pericellular domain [58, 59], to fusion and host-cell entry [60]. Each of these are complex processes involving a multitude of viral-host interactions. The omicron variant has, for example, been found to have accelerated cilia-dependent cellular entry in nasal epithelia which contributes to its increased attack rate over previous variants [60]. Many pre-omicron SARS-CoV-2 mutations are hypothesized to have increased fitness in viral attachment predominantly through enhanced ACE2 binding affinity achieved through spike and RBD-specific mutations [35, 37, 40, 41] (see Table 1). Some mutations, for example N439K, are found to enhance ACE2 binding and also contribute to humoral immune escape [61]. Although substantial increases in *α*, relative to viral clearance, are required to achieve the same *R*_0_ (Fig. 2B), we find the probability of transmitting mutants with higher attachment rates to be very similar to those reducing viral clearance, *C* (Fig. 3). Furthermore, the survival probability (Eq. S5) scales with bottleneck size similarly for virion attachment as it does for virion clearance (See Fig.S2). Therefore, our work suggests mutations affecting humoral immune evasion and ACE2 binding affinity and fusogenicity to be equally transmittable, assuming the mutations confer a similar selective advantage.

When an infectious virus enters its host cell it does not instantaneously begin self replication. Viral production requires a period of time, the eclipse phase, during which the virus manipulates a multitude of cellular processes to facilitate an environment for self replication and continued transmission [62]. In our model, advantageous mutations affecting the eclipse phase result in a virus more quickly transitioning from host-cell entry through to initial budding, captured through an increased eclipse rate, *E*. We find advantageous mutations affecting *E* to confer a high probability of transmission, Π. For example, an increase of 10% in *R*_0_ leads to an order of magnitude increase in Π, where this substantial increase in Π is not seen for any other parameter (Fig. 3). In Fig 4, showing Π as a function of bottleneck size, we predict that at a 0.5% *R*_0_ increase, favourable eclipse phase mutations have higher chances of transmission than equivalent changes to any other parameter, across any bottleneck size. As *R*_0_ is further increased to 5.0%, advantageous eclipse phase mutations lead to an order of magnitude increase in Π for all Λ, as compared to all other parameters. An alternative visualization, in which all the curves in Fig 4 are normalized by the neutral strain probability, is shown in Fig.S4. Here we see that at a 5.0% *R*_0_ increase (Fig.S4B), for narrow bottlenecks, Π exceeds 20 times the neutral strain value. As bottleneck size is increased, the neutralnormalized Π decreases monotonically such that at Λ = 50 Π has decreased to *∼*10 times the neutral strain value. Thus, we uncover that one outcome of increasing bottleneck size is to monotonically reduce the probability of transmission relative to the neutral strain. This effect is particularly apparent for mutations affecting the eclipse period, however, this trend can be observed with all other parameters as well (Fig S4). This result, which emerges from our analytical approach (Eq. S2), suggests that small bottleneck sizes [28, 29] favour the adaptive evolution of SARS-CoV-2, as we find narrow bottleneck sizes (Λ *<* 5) increase the transmission probability for adaptive, as compared to neutral, phenotypic changes.

One of the major components of the host-cell infection process is to reprogram cell metabolic processes, typically in the form of increased glycolysis [63] and hijacking of glucose transport pathways [64]. The SARS-CoV-2 M protein, the most abundant of the structural proteins [65], has been found to be homologous to the prokaryotic sugar transport protein SemiSWEET, and is further hypothesized to be implicated in proliferation, replication, and immune evasion [34]. Utilizing SARS-CoV-2 genomic surveillance data from 2020, with phylogenetic analyses carried out via Nextstrain [5], Shen *et al*. [66] associate the rapid emergence of the B.1.575 lineage with mutations in the M protein. They relate the mutations in the M protein to more advantageous glucose uptake during viral assembly [66]. In our model, this M mutation can be captured by advantageous mutations in the eclipse parameter, *E*, and/or the budding rate, *B*, as the mutation is hypothesized to lead to a more optimal within-cell replication and viral proliferation. In other modelling work, SARS-CoV-2 viral load has been shown to be highly sensitive to the eclipse duration; thus, mutations affecting the ability to more quickly reprogram intracellular machinery would be highly favourable [67]. It is therefore possible that our eclipse mutation results (Fig. 3), with neutral strain properties [23] determined from SARS-CoV-2 data gathered at a similar time as the Shen *et al*. [66] study, are revealing the high probability of such a mutation to emerge at this time of the pandemic, and this result further supports the importance of therapeutics that target the M protein [65].

SARS-CoV-2 mutations affecting immune escape have been of paramount interest throughout the pandemic [68, 69], and are of key importance towards understanding and predicting vaccine efficacy against emerging variants [70, 71]. Single mutations in M, E, N, or S can have complex outcomes with multiple phenotypic changes [61]. S mutations, for example, have been linked to enhanced viral entry (L5F), but also to increased resistance to antibodies (G446V and A879V) [9]. In our model, mutations enhancing immune escape could be captured by cell death, *D*, or viral load clearance, *C*. Mutations affecting *D* will include phenotypic outcomes associated with recognition or killing of infected cells, avoiding recognition by the immune system, or, if we extended the model to include a time-varying death rate, reducing the innate immune response. For example, the Alpha-variant mutations to Orf9b are associated with suppression of the innate immune response through reduced interferon induction [72]; such a change in viral dynamics could be captured by parameter *D* in our model. Advantageous mutations leading to humoral evasion are captured by decreases to the viral load clearance rate. For example, the BA.4 and BA.5 subvariants have been found to exhibit humoral evasion [73], and thus, would exhibit a lower rate of virion clearance as compared to previous mutants.

We find mutants affecting *D* to have little impact on predicted survival probabilities, (*P*_*s*_, Eq. S5), for all bottleneck sizes (Fig.S2D), and lead to transmission probabilities nearly equivalent to the neutral strain for all selective coefficients and bottleneck sizes we examined (Fig.S4, 4). In contrast, mutations in virion clearance scale similarly to target-cell attachment (Fig.S4, 4, S5C), and exhibit large increases in *P*_*s*_ as a function of increased bottleneck size for any particular *t*_*m*_ (Fig.S2C). This point is also made clear through a simple calculation of maximum predicted Π achieved for each parameter at narrow bottlenecks (Λ = 1). The percent difference in *D* required to achieve a 50% increase in *R*_0_ is 32% and results in an increase in Π of a factor of 1.3 relative to the neutral strain. In contrast, the percent difference in *C* required to achieve a 50% percent increase in *R*_0_ is 57% and results in an increase in Π of a factor of 15.1 relative to the neutral strain. Our results predict advantageous mutations leading to humoral escape have a significantly higher probability to be transmitted to the next host, as compared to mutations associated with recognition or killing of infected cells. However, we also note that if we restrict our results to an experimentally-estimated Π range of 10^−7^*−* 10^−6^ (grey region, Fig 3), then the entire explored parameter space for *D* falls within the region, while only a limited region of parameter space for *C* does. Thus, although mutations in *C* confer a significantly higher transmission advantage, there appears to be significantly more explorable parameter space for mutations affecting *D*.

## S1 Supplementary material

### S1.1 Wildtype population dynamics

While rare mutations will be modelled using a stochastic approach (see S1.3), we use a standard, deterministic approach to model the dynamics of the bulk of the within-host infection; the “wildtype” infection. The wildtype model tracks the density of susceptible target cells (*y*_*T*_), cells in an eclipse stage but not yet able to produce virus (*y*_*j*_), infected cells capable of viral budding (*y*_*B*_), infectious virus not yet attached to susceptible cells (*v*) and non-infectious virus (*w*). Target cells are infected via a per-target cell attachment rate, *α*. Susceptible cells then enter into the first eclipse stage, *y*_1_, and progress through a total of *k* eclipse stages at rate of *kE* before entering the budding stage. This chain of eclipse stages yields an Erlang-distributed eclipse duration with shape parameter *k*, which has been shown to yield significantly better fits to SARS-CoV-2 viral load data than a single eclipse-phase compartment, where *k* = 3 was found to produce the lowest AIC and BIC [23]. We assume both eclipse and budding cells have a constant loss rate, *D*. Virus is produced by infected cells at constant rate *B*, and fraction *ϵ* of this virus is capable of infecting susceptible target cells; the remaining 1 *− ϵ* is non-infectious virus. Similar to previous within-host SARS-CoV-2 modelling work [23, 24], we include a term for loss of infectious virus due to successful virus entry into susceptible target cells. The full deterministic model with *k* = 3 eclipse stages is shown in Eq. S1, a schematic of this model is provided in Fig. 1a.

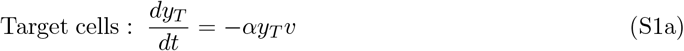

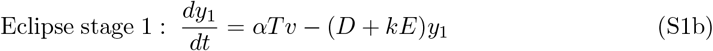

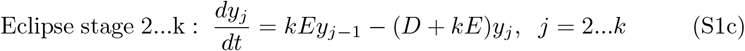

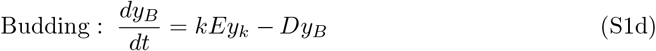

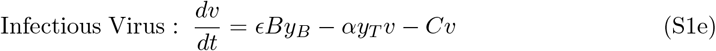

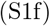

### S1.2 Parameter estimates and initial conditions

We use the parameter estimates from ref. [23], fitted to within-host data from ref. [21], to define the wildtype (WT) strain in the deterministic model, as these parameters were determined to have the best identifiability and agreed well with previous literature. Similar to ref.[20, 23, 24, 74], we assume an initial target cell concentration of 1.33×10^5^ cells*·*mL^*−*1^, and an initial condition of one productively infected cell in the upper respiratory tract, yielding *y*_*B*_(0) = 1 cell/30 mL. All other populations are initially zero. Parameter assumptions and initial conditions are summarized in Table S1.

**Table S1.** Parameter estimates for the wildtype (deterministic) model (Eq.1) and the mutant (stochastic) model (Eq.2). All parameters used in this work are SARS-CoV-2 specific, and sourced from previously published literature. † Total range explored in this work, motivated from all cited references.

| Parameter | Units | Definition | Estimate | Reference |
| --- | --- | --- | --- | --- |
| Deterministic model, parameters for the wildtype strain |  |  |  |  |
| $\alpha$ | mL·d<sup>-1</sup>·copies<sup>-1</sup> | Per target cell attachment rate | 7.9x10<sup>-5</sup> | [23] |
| $\epsilon$ | Unitless | Fraction of infectious virus | 10<sup>-4</sup> | [20, 25] |
| $\epsilon B$ | copiesd<sup>-1</sup>·d<sup>-1</sup>·cell<sup>-1</sup> | Infectious virus budding rate | 15.7 | [23] |
| $C$ | d<sup>-1</sup> | Clearance rate | 15 | [22, 23] |
| $D$ | d<sup>-1</sup> | Infected cell death rate | 0.32 | [23] |
| $E$ | d<sup>-1</sup> | Eclipse rate | 5 | [23, 75, 76] |
| $k$ | Unitless | Stages in eclipse phase | 3 | [23] |
| $R_0^N$ | Unitless | Reproduction number for the neutral strain | 19.0 | [23] |
| $y_T(0)$ | cell/mL | Initial number of target cells | 1.33x10<sup>5</sup> | [20, 24, 74] |
| $y_k(0)$ | cell/mL | Initial number of cells in eclipse phase | 0 | N/A |
| $y_B(0)$ | cell/mL | Initial number of infected cells | 1/30 | [20] |
| $v(0)$ | copies·mL<sup>-1</sup> | Initial number of infectious virus | 0 | N/A |
| Stochastic model parameters |  |  |  |  |
| $\Lambda$ | Virus particles | Bottleneck size: number of viruses founding infection | 1-50† | [9, 27–29] |
| $\mu$ | nt<sup>-1</sup>·cycle<sup>-1</sup> | Mutation rate (per site replication) | 1.3x10<sup>-6</sup> | [3] |
| $\tau$ | h | Disease transmission time | 144 | Calculated |

### S1.3 Stochastic model

To describe the fate of an initially rare SARS-CoV-2 mutation, we use a stochastic model previously developed for influenza A virus [31]. We consider single mutations (not chains of mutations), and assume that the mutant lineage propagates in an environment determined by the within-host model given by Eq. S1. The reproduction of the mutant lineage, as well as the corresponding eclipse stage cells and budding cells, is modelled as a branching process and described by the multitype probability generating function (pgf) *G*(*x*_1_, *x*_2_, …, *x*_*B*_, *t*). Here, dummy variables *x*_1_, *x*_*j*_, and *x*_*B*_ represent infectious viral load, eclipse stage cells, and budding cells, respectively. As derived previously [31], the time evolution of this pgf is described by

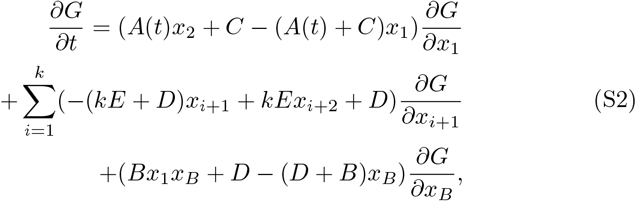

where all parameters except *A* are as described previously.

### S1.4 Model Coupling

The stochastic model describing the fate of rare mutant lineages is coupled to the deterministic model of the wildtype population in three ways. First, *A*(*t*) in Eq.S2 is the time-dependent per-virion attachment rate. We set this as *A*(*t*) = *αy*_*T*_ (*t*), where *y*_*T*_ (*t*) is the susceptible target cell density from the deterministic model, determined by solving Eq. S1 with parameter estimates in Table S1. We thus assume that the number of available target cells at any time is determined by the dynamics of the wildtype infection, and is not affected by rare mutations.

Secondly, we note that disease transmission can occur at any time in this model. For the numerical results to follow, we explore the case in which disease transmission occurs at time *τ*, the time of peak wildtype viral load. We thus use the deterministic model (Eq. S1) under the parameters in Table S1 to find the peak time *τ* = 144 h. In addition, the size of the transmission bottleneck (the number of virions that are transmitted to the next host) is given by Λ. Previous experimental estimates in SARS-CoV-2 have found Λ*≈* 1-3 [28] and Λ *≈* 2-47 [27]. We therefore examine a range of values from 1 to 50. Note that the free virus in Eq. S1 is modelled as a density in a 30 mL volume of the upper respiratory tract. Thus the number of viral particles at any time is given by 30*v*(*t*). We assume each viral particle is equally likely to survive the transmission bottleneck and be passed on to the next host. We thus define the *F*, the probability that a virion survives the bottleneck, as

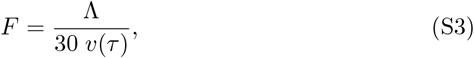

since *v*(*τ*) is the concentration of viral load at transmission. The probability *F* is therefore determined by the wildtype infection.

Thirdly, rare mutations can occur at any time during the infection. We assume that each replication of a virion is subject to mutation rate *μ* (per site per replication), and thus the density of new mutant lineages that first occur at time *t*_*m*_ is given by *μBy*_*B*_(*t*_*m*_), where again these values are determined by the wildtype infection. We take *μ* = 1.3 *×*10^−6^, as recently estimated for SARS-CoV-2 [3], and we neglect the chance of further mutation to these rare mutant lineages (but see [30]).

To solve Eq. S2 we use the method of characteristics to produce a system of ordinary equationsn (Eq. S9) which are then implemented and solved numerically in **R** (version 2022.12.0+353).

### S1.5 Selective effects of mutations

The stochastic model (Eq. S2), includes the life-history parameters of the virus, *α, B, C* etc. When these parameters are set to the wildtype values (Table S1), the stochastic model tracks the fate of rare mutations that are neutral, that is, they have no phenotypic effect. In addition to quantifying the fate of neutral mutations, the goal of this work is to predict the fate of *de novo* mutations that confer an adaptive advantage to the virus. To compare parameter values that give an equivalent adaptive advantage, we use the within-host basic reproductive ratio, *R*_0_. (We emphasize that this within-host *R*_0_ reflects the number of infectious virions produced on average by a single infectious virion, and does not correspond to the epidemiological *R*_0_ for SARS-CoV-2 [77]). For Eq. S1 with *k* eclipse stages, an expression for *R*_0_ is derived in ref. [23] and is given by:

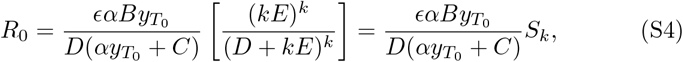

where *S*_*k*_ is the probability that an infected cell survives all *k* eclipse stages [23]. For Table S1 parameters, this yields a value of 19.0 for wildtype and neutral variants, 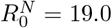.

To study the fate of an adaptive *de novo* mutation, we assume that the mutation changes one life-history trait of the virus, leading to an increase in *R*_0_. We consider single mutations that lead to an increase in *R*_0_ from 0.5% to 50%. For parameters involved in immune escape, trait changes are modelled using reductions in the viral clearance or cell death rate: 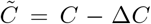 or 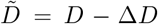, respectively. Beneficial mutations leading to an increase in viral attachment, budding or infected cell maturation rates are described using parameters 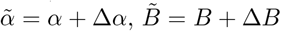 or 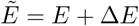, respectively.

### S1.6 Mutant lineage transmission probability

Each infection begins with the founding virus of the wildtype strain at time *t* = 0*h* as described for the wildtype model in Section S1.1. At time *t* = *t*_*m*_, such that 0 *≤t*_*m*_ *≤ τ*, a *de novo* mutation, either neutral or with a selective advantage, appears. To determine the transmission probability of a particular mutant lineage that first arose at time *t*_*m*_, we solve Eq. S2 (as described in ref. [31]). In particular, we use the initial condition corresponding to a single free virus at time *t*_*m*_, and we replace any life-history parameters that have changed in the mutant strain by their corresponding value.

The solution of Eq. S2 at time *τ* gives the distribution of the mutant lineage just before transmission, given the mutated lineage began with a single virion at time *t*_*m*_. To determine the probability, *χ*(*t*_*m*_), that a particular mutant lineage does not survive transmission at time *τ*, we take advantage of the well-known properties of pgfs and simply evaluate *G* at *x*_1_ = 1 *− F, x*_2_ = *x*_3_ = 1, *t* = *τ* (see [31] for details). We then write the ‘survival probability’, *P*_s_, as:

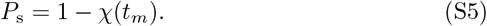

In summary, *P*_s_ is the probability that at least one viral particle is transmitted to the next host, from a mutant lineage that arose as a single virion at time *t*_*m*_.

Given this survival probability, we then compute the rate, at any time, at which new mutant lineages that will survive the bottleneck event emerges:

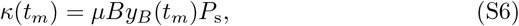

where *μBy*_*B*_(*t*_*m*_) is the rate at which new mutant lineages are created, as described previously [31]. We can then integrate *κ* to compute, *S*, the expected number of times, during a single infection, that a mutant emerges that will survive to be transmitted:

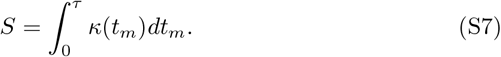

This allows us to compute Π, defined as the probability that the mutation arises during the infection, and at least one copy survives to be transmitted

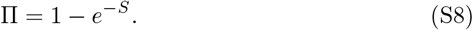

The quantity Π thus estimates the probability that a new variant emerges and is passed on during a single infection.

### S1.7 Method of characteristics

The time evolution of the pgf given by Eq.2 describes the fate of an initially rare SARS-CoV-2 mutation. The method of characteristics is used on Eq.2 to produce a system of ordinary differential equations (Eq.S1), which is then implemented and solved in **R**.

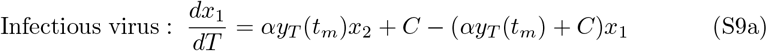

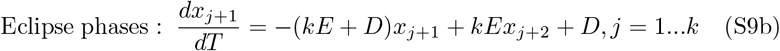

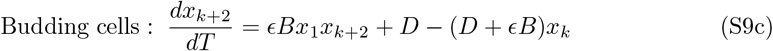

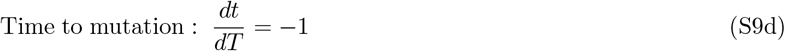

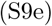

### S1.8 Narrow bottleneck (Λ = 1) results

#### S1.8.1 Neutral strain results and mutant strain results

**Fig. S1.**
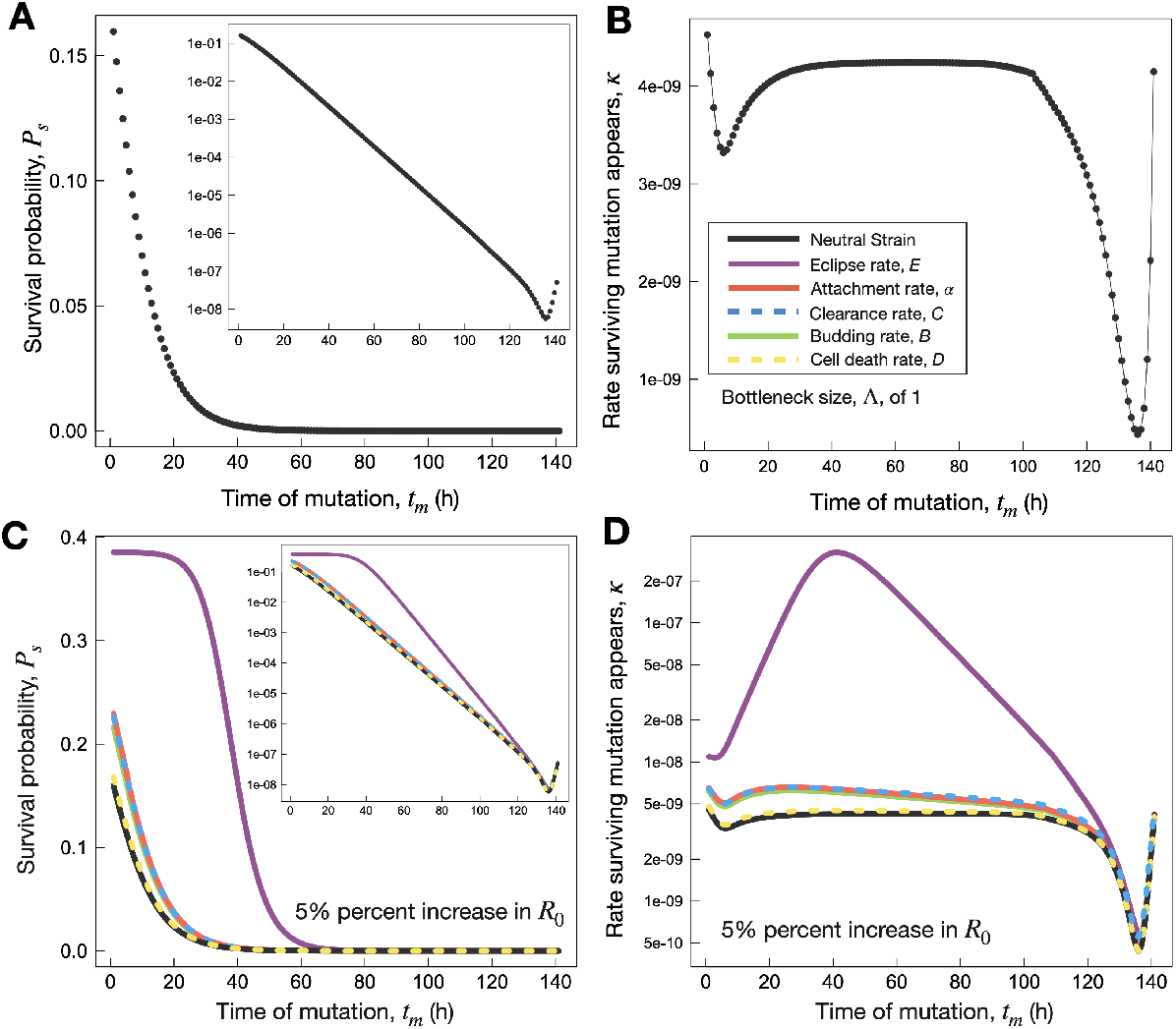
A) Survival probability, *P*_*s*_ (Eq. S5) as a function of time of mutation, *t*_*m*_. B) Rate surviving mutations appear, *κ* (Eq. S6), as a function of time of mutation, *t*_*m*_. C-D) Differences in parameters dynamics for Λ = 1 and a fixed selective coefficient of 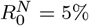. C) Survival probability, *P*_*s*_ (Eq. S5), as a function of *t*_*m*_. D) Survival probability, *P*_*s*_ (Eq. S5), as a function of *t*_*m*_. The eclipse rate, *E*, is observed to have a more pronounced affect whereby *P*_*s*_ is as high as 0.38 compared to *P*_*s*_ *≈* 0.2 for all other parameters at *t*_*m*_ *≈*0. Similarly, the peak in *κ* exceeds 2×10^−7^ for mutations in *E* leading to a 5% increase in 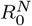, while all other parameters peak below 1×10^−8^.

#### S1.8.2 Survival probabilities and rate of appearance of surviving mutations

**Fig. S2.**
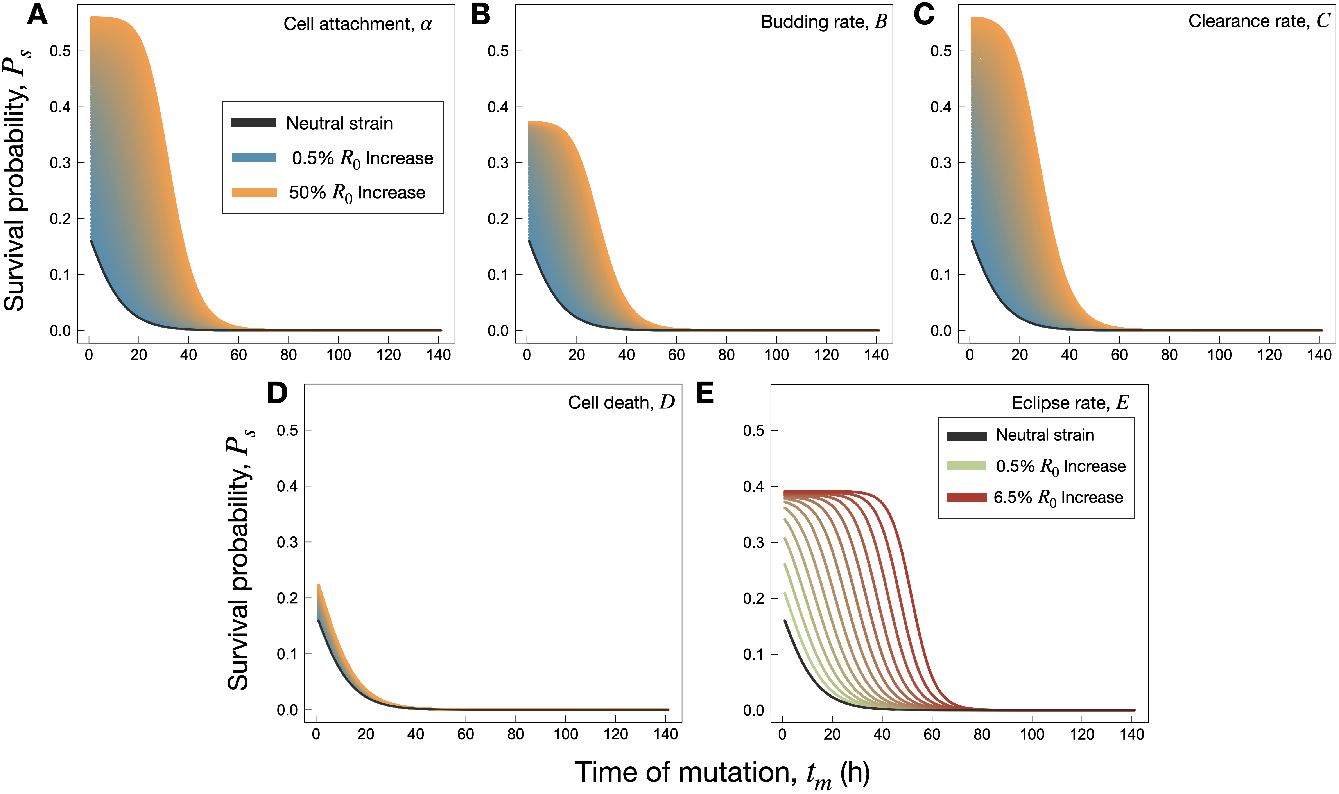
Survival probabilities, *P*_*s*_, for narrow bottlenecks of size Λ = 1. A-E) *P*_*s*_ as a function of *t*_*m*_ for *α, B, C, D*, and *E*, respectively. Note *E* is limited to a maximum selective coefficient of 6.5%, and is therefore shown with a different colour scheme. Mutations in *α* and *C* are found to increase *P*_*s*_ by a similar amount for each 0.5% increase in 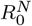, whereas mutations affecting *B* are intermediate, and mutations affecting *D* lead to a negligible increase in *P*_*s*_. Mutations in *E* are found to increase *P*_*s*_ from the neutral strain up to *∼* 0.4, further, mutations in *E* lead to *P*_*s*_ maintaining maximum value as a function of *t*_*m*_ for an extended period of time, as compared to mutations in the other parameters.

**Fig. S3.**
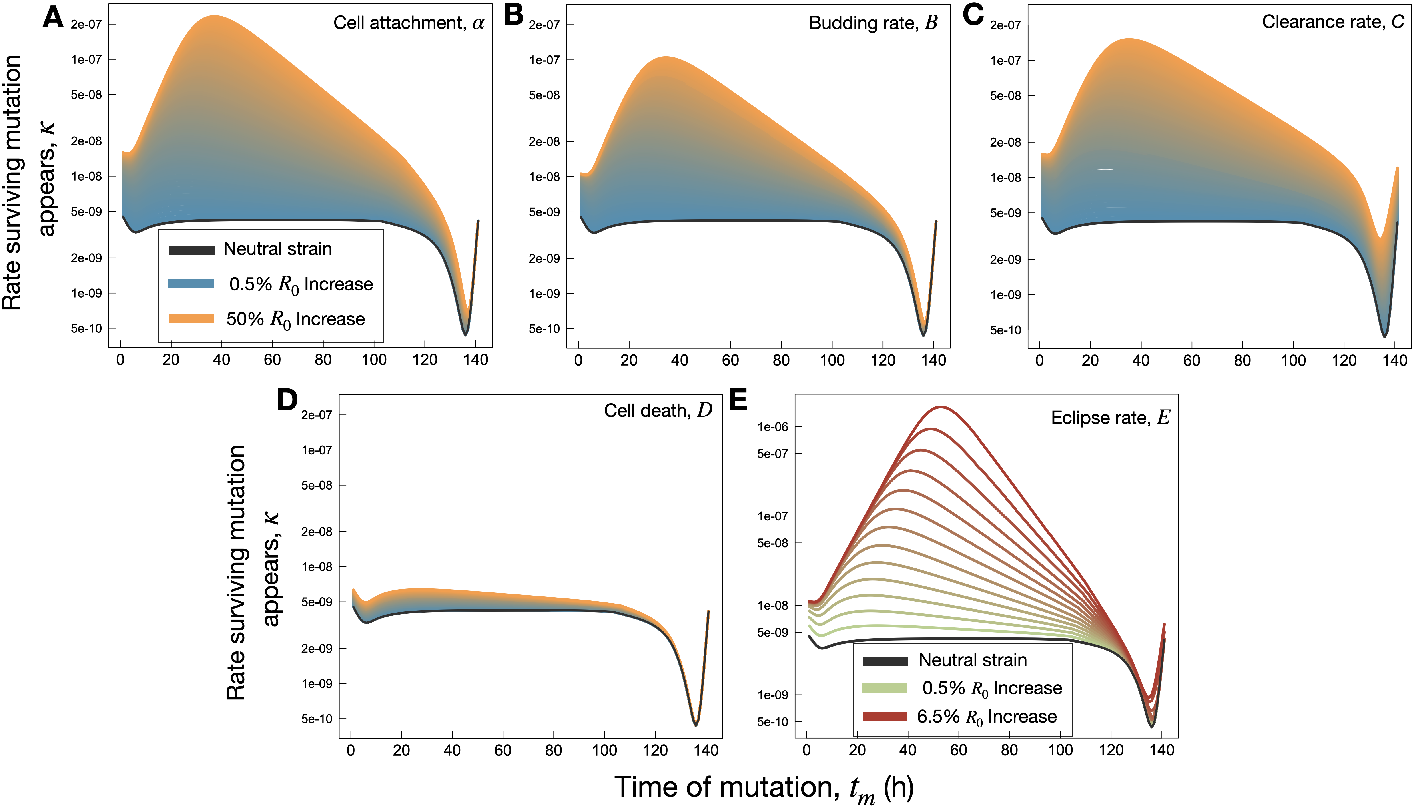
Rate of appearance of surviving mutations, *κ*, for narrow bottlenecks of size Λ = 1. A-E) *κ* as a function of *t*_*m*_ for *α, B, C, D*, and *E*, respectively. Note *E* is limited to a maximum selective coefficient of 6.5%, and is therefore shown with a different colour scheme. Mutations in *α, B*, and *C* all have a pronounced effect on *κ*, whereby the formation of a peak value at *t*_*m*_ = 40h emerges. Mutations in *D* leading to a 50% increase in 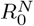 have little affect on *κ*. Mutations in *E* lead to the formation of a peak in *κ* that shifts towards later values as 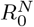 is increased from 0.5% to 6.5%.

#### S1.9 Mutation dynamics for increasing Λ

To further illustrate the selective advantage of each mutation over the neutral strain, in Fig.S4 we report the ratio of the mutant Π to that of the neutral strain for each parameter. Here, we see that at 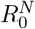 increase of 0.5% the maximum advantage in Π over the neutral strain is 1.25x higher for *E*, whereby all other parameters are under 1.05x higher transmission probability over the neutral strain. At a selective coefficient of 5.0% *E* has a 20x higher probability of transmitting at narrow bottlenecks, while at large bottlenecks of Λ = 50 the advantage reduces to 10x, as compared to the neutral strain. In comparison, all other parameters are below 1.5x relative to the neutral strain, while Π for mutations in *D* for all Λ are approximately equal to Π of the neutral strain. When the selective coefficient is 50% *α* and *B* have a 20x higher probability of transmitting at narrow bottlenecks,while at large bottlenecks of Λ = 50 the advantage reduces to 10x, as compared to the neutral strain. Whereas mutations in *B* lead to a 10x higher probability of transmission at narrow bottlenecks and *∼*5x higher probability at large bottlenecks. Mutations in *D*, however, maintain a much smaller advantage in transmitting at less than 1.5x more than the neutral strain for all Λ.

**Fig. S4.**
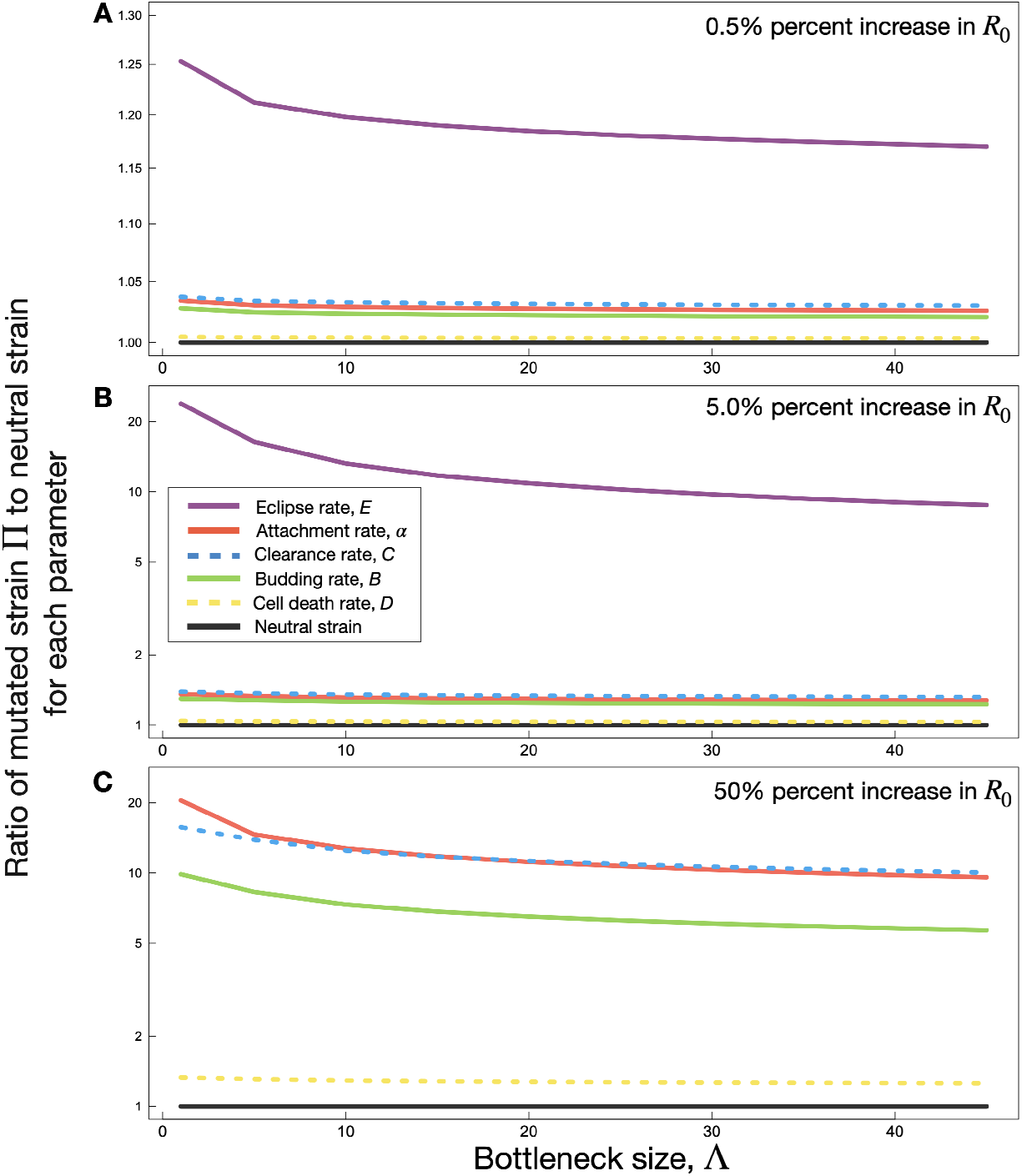
A-C) Ratio of Probability of transmitting at least one mutant, Π (Eq. S8), to that of the neutral strain, as a function of bottleneck size, Λ, for selective coefficients 0.5% (panel A), 5% (panel B), and 50% (panel C). *E* is omitted from panel C as only selective coefficients up to 6.5% were examined for this parameter.

Fig. S5 examines these trends in greater detail, combing results for Π as a function of both *R*_0_ and bottleneck size. Fig. S5 panels A-E correspond to mutations affecting parameters *α, B, C, D*, and *E*, respectively, while the gradient colour code of blue to orange corresponds to Λ = 1 to Λ = 50 as indicated. In all cases, Π increases monotonically with both the fitness of the mutant and with bottleneck size. Noting the log-scale of the *y*-axis, we see that the increase in Π is roughly exponential with mutant fitness, for mutations affecting *α, B, C* and *D*, although the rate of increase in the latter case is small. We also find that Π scales faster than exponentially as a function of *R*_0_ for mutations affecting *E*.

**Fig. S5.**
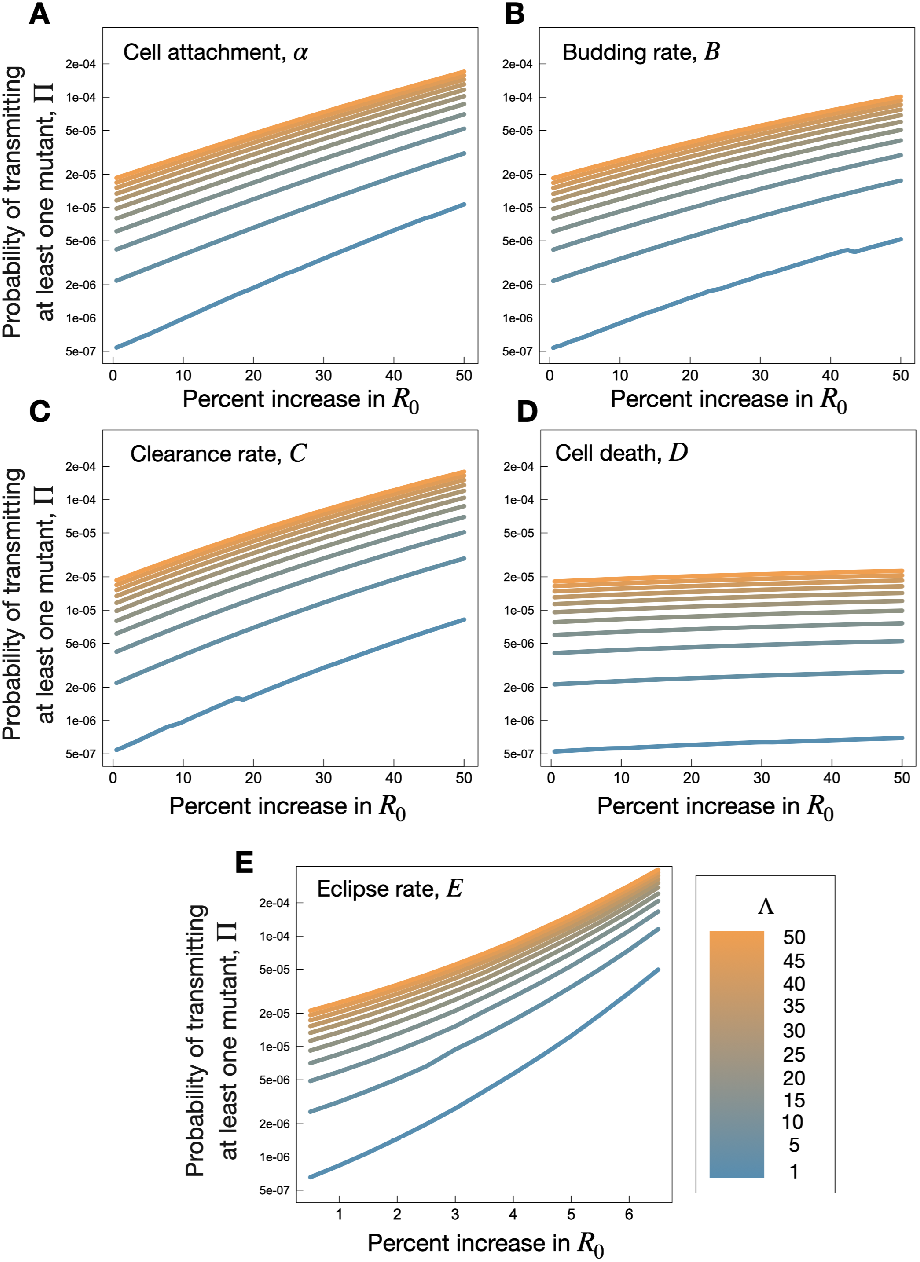
The probability of transmitting at least one mutant, Π, as a function of the percent increase in *R*_0_. For each parameter, the bottleneck size is illustrated with a colour gradient from blue (Λ = 1) to orange (Λ = 50).

## Notes

### Competing Interest Statement

The authors have declared no competing interest.

## References

[1] Rambaut, A., Holmes, E.C., O’Toole, Á., Hill, V., McCrone, J.T., Ruis, C., du Plessis, L., Pybus, O.G.: A dynamic nomenclature proposal for SARS-CoV-2 lineages to assist genomic epidemiology. Nature Microbiology 5(11), 1403–1407 (2020). https://doi.org/10.1038/s41564-020-0770-5

[2] XBB.1.5 Updated Risk Assessment, 24 February 2023. World Health Organization (2023)

[3] Amicone, M., Borges, V., Alves, M.J., Isidro, J., Zé-Zé, L., Duarte, S., Vieira, L., Guiomar, R., Gomes, J.P., Gordo, I.: Mutation rate of SARS-CoV-2 and emergence of mutators during experimental evolution. Evolution, Medicine and Public Health 10(1), 142–155 (2022). https://doi.org/10.1093/emph/eoac010

[4] HO Coronavirus (COVID-19) Dashboard. World Health Organization (2023)

[5] Hadfield, J., Megill, C., Bell, S.M., Huddleston, J., Potter, B., Callender, C., Sagulenko, P., Bedford, T., Neher, R.A.: Nextstrain: real-time tracking of pathogen evolution. Bioinformatics 34(23), 4121–4123 (2018)

[6] Zhang, C., Gu, J., Chen, Q., Deng, N., Li, J., Huang, L., Zhou, X.: Clinical and epidemiological characteristics of pediatric SARS-CoV-2 infections in China: A multicenter case series. PLoS medicine 17(6), 1003130 (2020)

[7] Salzberger, B., Buder, F., Lampl, B., Ehrenstein, B., Hitzenbichler, F., Holzmann, T., Schmidt, B., Hanses, F.: Epidemiology of SARS-CoV-2. Infection 49, 233–239 (2021)

[8] Tian, D., Sun, Y., Xu, H., Ye, Q.: The emergence and epidemic characteristics of the highly mutated SARS-CoV-2 Omicron variant. Journal of medical virology 94(6), 2376–2383 (2022)

[9] Lythgoe, K.A., Hall, M., Ferretti, L., de Cesare, M., MacIntyre-Cockett, G., Trebes, A., Andersson, M., Otecko, N., Wise, E.L., Moore, N., Lynch, J., Kidd, S., Cortes, N., Mori, M., Williams, R., Vernet, G., Justice, A., Green, A., Nicholls, S.M., Ansari, M.A., Abeler-Dörner, L., Moore, C.E., Peto, T.E.A., Eyre, D.W., Shaw, R., Simmonds, P., Buck, D., Todd, J.A., Connor, T.R., Ashraf, S., da Silva Filipe, A., Shepherd, J., Thomson, E.C., Bonsall, D., Fraser, C., Golubchik, T.: SARS-CoV-2 within-host diversity and transmission. Science 372(6539) (2021). https://doi.org/10.1126/SCIENCE.ABG0821

[10] Gholami, S., Korosec, C.S., Farhang-Sardroodi, S., Dick, D.W., Craig, M., Ghaemi, M.S., Ooi, H.K., Heffernan, J.M.: A mathematical model of protein subunits COVID-19 vaccines. Mathematical Biosciences 358, 108970 (2023). https://doi.org/10.1016/J.MBS.2023.108970

[11] Padmanabhan, P., Desikan, R., Dixit, N.M.: Modeling how antibody responses may determine the efficacy of COVID-19 vaccines. Nature Computational Science 2(2), 123–131 (2022). https://doi.org/10.1038/s43588-022-00198-0

[12] Lin, J., Law, R., Korosec, C.S., Zhou, C., Koh, H., Ghaemi, S., Samaan, P., Kiang Ooi, H., Matveev, V., Yue, F., Gingras, A.-C., Estacio, A., Buchholz, M., Cheatley, P.L., Mohammadi, A., Kaul, R., Pavinski, K., Mubareka, S., McGeer, A.J., Leis, J.A., Heffernan, J.M., Ostrowski, M.: Longitudinal Assessment of SARS-CoV-2-Specific T Cell Cytokine-Producing Responses for 1 Year Reveals Persistence of Multicytokine Proliferative Responses, with Greater Immunity Associated with Disease Severity. Journal of Virology 96(13), 00509–22 (2022). https://doi.org/26

[13] Korosec, C.S., Farhang-Sardroodi, S., Dick, D.W., Gholami, S., Ghaemi, M.S., Moyles, I.R., Craig, M., Ooi, H.K., Heffernan, J.M.: Long-term durability of immune responses to the BNT162b2 and mRNA-1273 vaccines based on dosage, age and sex. Scientific Reports 12(1), 21232 (2022) arXiv:/doi.org/10.1101/2021.10.13.21264957 [https:]. https://doi.org/10.1038/s41598-022-25134-0

[14] Moyles, I.R., Korosec, C.S., Heffernan, J.M.: Determination of significant immunological timescales from mRNA-LNP-based vaccines in humans. Journal of Mathematical Biology 86(86), 1–41 (2023) arXiv:2022.07.25.22278031. https://doi.org/10.1007/s00285-023-01919-3

[15] Farhang-sardroodi, S., Korosec, C.S., Gholami, S., Craig, M., Moyles, I.R., Ghaemi, M.S., Ooi, H.K., Heffernan, J.M.: Analysis of Host Immunological Response of Adenovirus-Based COVID-19 Vaccines. Vaccines 9(8), 861 (2021)

[16] Matveev, V.A., Mihelic, E.Z., Benko, E., Budylowski, P., Grocott, S., Lee, T., Korosec, C.S., Colwill, K., Stephenson, H., Law, R., Ward, L.A., Sheikh-Mohamed, S., Mailhot, G., Delgado-Brand, M., Pasculescu, A., Wang, J.H., Qi, F., Tursun, T., Kardava, L., Chau, S., Samaan, P., Imran, A., Dennis C. Copertino, J., Chao, G., Choi, Y., Reinhard, R.J., Kaul, R., Heffernan, J.M., Jones, R.B., Chun, T.-W., Moir, S., Singer, J., Gommerman, J., Gingras, A.-C., Kovacs, C., Ostrowski, M.: Immunogenicity of COVID-19 vaccines and their effect on the HIV reservoir in older people with HIV. bioRxiv, 2023–0614544834 (2023). https://doi.org/10.1101/2023.06.14.544834

[17] Betti, M., Bragazzi, N.L., Heffernan, J.M., Kong, J., Raad, A.: Integrated vaccination and non-pharmaceutical interventions based strategies in Ontario, Canada, as a case study: a mathematical modelling study. Journal of The Royal Society Interface 18(180), 20210009 (2021)

[18] Johansson, M.A., Quandelacy, T.M., Kada, S., Prasad, P.V., Steele, M., Brooks, J.T., Slayton, R.B., Biggerstaff, M., Butler, J.C.: SARS-CoV-2 transmission from people without COVID-19 symptoms. JAMA network open 4(1), 2035057–2035057 (2021)

[19] Le Rutte, E.A., Shattock, A.J., Chitnis, N., Kelly, S.L., Penny, M.A.: Modelling the impact of Omicron and emerging variants on SARS-CoV-2 transmission and public health burden. Communications Medicine 2(1), 93 (2022)

[20] Néant, N., Lingas, G., Le Hingrat, Q., Ghosn, J., Engelmann, I., Lepiller, Q., Gaymard, A., Ferré, V., Hartard, C., Plantier, J.-C., Thibault, V., Marlet, J., Montes, B., Bouiller, K., Lescure, F.-X., Timsit, J.-F., Faure, E., Poissy, J., Chidiac, C., Raffi, F., Kimmoun, A., Etienne, M., Richard, J.-C., Tattevin bb, P., Garot cc, D., Le Moing dd, V., Bachelet ee, D., Tardivon ee, C., Duval, X., Yazdanpanah, Y., Mentré, F., Laouénan, C., Visseaux, B., Guedj, J.: Modeling SARS-CoV-2 viral kinetics and association with mortality in hospitalized patients from the French COVID cohort. PNAS 118(8), 2017962118 (2021). https://doi.org/10.1073/pnas.2017962118/-/DCSupplemental

[21] Jones, T.C., Biele, G., Mühlemann, B., Veith, T., Schneider, J., Beheim-Schwarzbach, J., Bleicker, T., Tesch, J., Schmidt, M.L., Sander, L.E., Kurth, F., Menzel, P., Schwarzer, R., Zuchowski, M., Hofmann, J., Krumbholz, A., Stein, A., Edelmann, A., Corman, V.M., Drosten, C.: Estimating infectiousness throughout SARS-CoV-2 infection course. Science 373(6551), 5273 (2021). https://doi.org/10.1126/science.abi5273

[22] Goyal, A., Cardozo-Ojeda, E.F., Schiffer, J.T.: Potency and timing of antiviral therapy as determinants of duration of SARS-CoV-2 shedding and intensity of inflammatory response. Science Advances 6(47), 7112 (2020). https://doi.org/10.1126/sciadv.abc7112

[23] Korosec, C.S., Betti, M.I., David, W., Ooi, H.K., Moyles, I.R., Wahl, L.M., Heffernan, J.M.: Multiple cohort study of hospitalized SARS-CoV-2 in-host infection dynamics: Parameter estimates, identifiability, sensitivity and the eclipse phase profile. Journal of theoretical biology 564, 111449 (2023) arXiv:2022.06.20.22276662. https://doi.org/10.1016/j.jtbi.2023.111449

[24] Gonçalves, A., Bertrand, J., Ke, R., Comets, E., de Lamballerie, X., Malvy, D., Pizzorno, A., Terrier, O., Rosa Calatrava, M., Mentré, F., Smith, P., Perelson, A.S., Guedj, J.: Timing of Antiviral Treatment Initiation is Critical to Reduce SARS-CoV-2 Viral Load. CPT: Pharmacometrics and Systems Pharmacology 9(9), 509–514 (2020). https://doi.org/10.1002/psp4.12543

[25] Gonçalves, A., Maisonnasse, P., Donati, F., Albert, M., Behillil, S., Contreras, V., Naninck, T., Marlin, R., Solas, C., Pizzorno, A., Lemaitre, J., Kahlaoui, N., Terrier, O., Fang, R.H.T., Enouf, V., Dereuddre-Bosquet, N., Brisebarre, A., Touret, F., Chapon, C., Hoen, B., Lina, B., Calatrava, M.R., de Lamballerie, X., Mentré, F., Le Grand, R., van der Werf, S., Guedj, J.: SARS-CoV-2 viral dynamics in non-human primates. PLoS Computational Biology 17(3), 1008785 (2021). https://doi.org/10.1371/JOURNAL.PCBI.1008785

[26] Perelson, A.S., Ke, R.: Mechanistic Modeling of SARS-CoV-2 and Other Infectious Diseases and the Effects of Therapeutics. Clinical Pharmacology and Therapeutics 109(4), 829–840 (2021). https://doi.org/10.1002/cpt.2160

[27] Braun, K.M., Moreno, G.K., Wagner, C., Accola, M.A., Rehrauer, W.M., Baker, D.A., Koelle, K., O’Connor, D.H., Bedford, T., Friedrich, T.C., Moncla, L.H.: Acute SARS-CoV-2 infections harbor limited within-host diversity and transmit via tight transmission bottlenecks. PLoS Pathogens 17(8), 1–26 (2021). https://doi.org/10.1371/journal.ppat.1009849

[28] Martin, M.A., Koelle, K.: Comment on “Genomic epidemiology of superspreading events in Austria reveals mutational dynamics and transmission properties of SARS-CoV-2”. Science Translational Medicine 13(617), 1803 (2023). https://doi.org/10.1126/scitranslmed.abh1803

[29] Bendall, E.E., Callear, A.P., Getz, A., Goforth, K., Edwards, D., Monto, A.S., Martin, E.T., Lauring, A.S.: Rapid transmission and tight bottlenecks constrain the evolution of highly transmissible SARS-CoV-2 variants. Nature Communications 14(1), 1–7 (2023). https://doi.org/10.1038/s41467-023-36001-5

[30] Van Egeren, D., Novokhodko, A., Stoddard, M., Tran, U., Zetter, B., Rogers, M.S., Joseph-McCarthy, D., Chakravarty, A.: Controlling longterm SARS-CoV-2 infections can slow viral evolution and reduce the risk of treatment failure. Scientific Reports 11(1), 1–9 (2021). https://doi.org/10.1038/s41598-021-02148-8

[31] Sigal, D., Reid, J.N.S., Wahl, L.M.: Effects of transmission bottlenecks on the diversity of influenza a virus. Genetics 210(3), 1075–1088 (2018). https://doi.org/10.1534/genetics.118.301510

[32] Sette, A., Crotty, S.: Adaptive immunity to SARS-CoV-2 and COVID-19. Cell 184(4), 861–880 (2021). https://doi.org/10.1016/j.cell.2021.01.007

[33] Longdon, B., Hadfield, J.D., Day, J.P., Smith, S.C.L., McGonigle, J.E., Cogni, R., Cao, C., Jiggins, F.M.: The Causes and Consequences of Changes in Virulence following Pathogen Host Shifts. PLoS Pathogens 11(3), 1–18 (2015). https://doi.org/10.1371/journal.ppat.1004728

[34] Thomas, S.: The Structure of the Membrane Protein of SARS-CoV-2 Resembles the Sugar Transporter SemiSWEET. Pathogens and Immunity 5(1), 342–363 (2020). https://doi.org/10.20411/pai.v5i1.377

[35] Liu, Y., Liu, J., Plante, K.S., Plante, J.A., Xie, X., Zhang, X., Ku, Z., An, Z., Scharton, D., Schindewolf, C., Widen, S.G., Menachery, V.D., Shi, P.-Y., Weaver, S.C.: The N501Y spike substitution enhances SARS-CoV-2 infection and transmission. Nature 602(7896), 294–299 (2022). https://doi.org/10.1038/s41586-021-04245-0

[36] Plante, J.A., Liu, Y., Liu, J., Xia, H., Johnson, B.A., Lokugamage, K.G., Zhang, X., Muruato, A.E., Zou, J., Fontes-Garfias, C.R., Mirchandani, D., Scharton, D., Bilello, J.P., Ku, Z., An, Z., Kalveram, B., Freiberg, A.N., Menachery, V.D., Xie, X., Plante, K.S., Weaver, S.C., Shi, P.-Y.: Spike mutation D614G alters SARS-CoV-2 fitness. Nature 592(7852), 116–121 (2021). https://doi.org/10.1038/s41586-020-2895-3

[37] Ozono, S., Zhang, Y., Ode, H., Sano, K., Tan, T.S., Imai, K., Miyoshi, K., Kishigami, S., Ueno, T., Iwatani, Y.: SARS-CoV-2 D614G spike mutation increases entry efficiency with enhanced ACE2-binding affinity. Nature communications 12(1), 848 (2021)

[38] Jangra, S., Ye, C., Rathnasinghe, R., Stadlbauer, D., Alshammary, H., Amoako, A.A., Awawda, M.H., Beach, K.F., Bermúdez-González, M.C., Chernet, R.L., Eaker, L.Q., Ferreri, E.D., Floda, D.L., Gleason, C.R., Kleiner, G., Jurczyszak, D., Matthews, J.C., Mendez, W.A., Mulder, L.C.F., Russo, K.T., Salimbangon, A.-B.T., Saksena, M., Shin, A.S., Sominsky, L.A., Srivastava, K., Krammer, F., Simon, V., Martinez-Sobrido, L., García-Sastre, A., Schotsaert, M.: SARS-CoV-2 spike E484K mutation reduces antibody neutralisation. The Lancet Microbe 2(7), 283–284 (2021). https://doi.org/10.1016/S2666-5247(21)00068-9

[39] Alenquer, M., Ferreira, F., Lousa, D., Valério, M., Medina-Lopes, M., Bergman, M.-L., Gonçalves, J., Demengeot, J., Leite, R.B., Lilue, J., Ning, Z., Penha-Gonçalves, C., Soares, H., Soares, C.M., Amorim, M.J.: Signatures in SARS-CoV-2 spike protein conferring escape to neutralizing antibodies. PLOS Pathogens 17(8), 1009772 (2021)

[40] Fratev, F.: N501Y and K417N mutations in the spike protein of SARS-CoV-2 alter the interactions with both hACE2 and human-derived antibody: a free energy of perturbation retrospective study. Journal of Chemical Information and Modeling 61(12), 6079–6084 (2021)

[41] Khan, A., Zia, T., Suleman, M., Khan, T., Ali, S.S., Abbasi, A.A., Mohammad, A., Wei, D.: Higher infectivity of the SARS-CoV-2 new variants is associated with K417N/T, E484K, and N501Y mutants: an insight from structural data. Journal of cellular physiology 236(10), 7045–7057 (2021)

[42] Tchesnokova, V., Kulasekara, H., Larson, L., Bowers, V., Rechkina, E., Kisiela, D., Sledneva, Y., Choudhury, D., Maslova, I., Deng, K.: Acquisition of the L452R mutation in the ACE2-binding interface of Spike protein triggers recent massive expansion of SARS-Cov-2 variants. Journal of clinical microbiology 59(11), 00921–21 (2021)

[43] Zhang, Y., Zhang, T., Fang, Y., Liu, J., Ye, Q., Ding, L.: SARS-CoV-2 spike L452R mutation increases Omicron variant fusogenicity and infectivity as well as host glycolysis. Signal Transduction and Targeted Therapy 7(1), 76 (2022). https://doi.org/10.1038/s41392-022-00941-z

[44] Motozono, C., Toyoda, M., Zahradnik, J., Saito, A., Nasser, H., Tan, T.S., Ngare, I., Kimura, I., Uriu, K., Kosugi, Y.: SARS-CoV-2 spike L452R variant evades cellular immunity and increases infectivity. Cell host & microbe 29(7), 1124–1136 (2021)

[45] Di Giacomo, S., Mercatelli, D., Rakhimov, A., Giorgi, F.M.: Preliminary report on severe acute respiratory syndrome coronavirus 2 (SARS-CoV-2) Spike mutation T478K. Journal of Medical Virology 93(9), 5638–5643 (2021). https://doi.org/10.1002/jmv.27062

[46] Wilhelm, A., Toptan, T., Pallas, C., Wolf, T., Goetsch, U., Gottschalk, R., Vehreschild, M.J.G.T., Ciesek, S., Widera, M.: Antibody-mediated neutralization of authentic SARS-CoV-2 B. 1.617 variants harboring L452R and T478K/E484Q. Viruses 13(9), 1693 (2021)

[47] Shen, L., Bard, J.D., Triche, T.J., Judkins, A.R., Biegel, J.A., Gai, X.: Emerging variants of concern in SARS-CoV-2 membrane protein: a highly conserved target with potential pathological and therapeutic implications. Emerging Microbes & Infections 10(1), 885–893 (2021). https://doi.org/10.1080/22221751.2021.1922097

[48] Lista, M.J., Winstone, H., Wilson, H.D., Dyer, A., Pickering, S., Galao, R.P., De Lorenzo, G., Cowton, V.M., Furnon, W., Suarez, N.: The P681H mutation in the spike glycoprotein of the alpha variant of SARS-CoV-2 escapes IFITM restriction and is necessary for type I interferon resistance. Journal of virology 96(23), 01250–22 (2022)

[49] Nie, C., Sahoo, A.K., Netz, R.R., Herrmann, A., Ballauff, M., Haag, R.: Charge matters: Mutations in omicron variant favor binding to cells. ChemBioChem 23(6), 202100681 (2022)

[50] Bills, C.J., Xia, H., Chen, J.Y.-C., Yeung, J., Kalveram, B., Walker, D., Xie, X., Shi, P.-Y.: Mutations in SARS-CoV-2 variant nsp6 enhance type-I interferon antagonism. Emerging Microbes & Infections (just-accepted), 2209208 (2023)

[51] Xia, S., Wang, L., Zhu, Y., Lu, L., Jiang, S.: Origin, virological features, immune evasion and intervention of SARS-CoV-2 Omicron sublineages. Signal Transduction and Targeted Therapy 7(1), 241 (2022). https://doi.org/10.1038/s41392-022-01105-9

[52] Fan, Y., Li, X., Zhang, L., Wan, S., Zhang, L., Zhou, F.: SARS-CoV-2 Omicron variant: recent progress and future perspectives. Signal transduction and targeted therapy 7(1), 141 (2022)

[53] Escalera, A., Gonzalez-Reiche, A.S., Aslam, S., Mena, I., Laporte, M., Pearl, R.L., Fossati, A., Rathnasinghe, R., Alshammary, H., van de Guchte, A.: Mutations in SARS-CoV-2 variants of concern link to increased spike cleavage and virus transmission. Cell host & microbe 30(3), 373–387 (2022)

[54] Saito, A., Irie, T., Suzuki, R., Maemura, T., Nasser, H., Uriu, K., Kosugi, Y., Shirakawa, K., Sadamasu, K., Kimura, I.: Enhanced fusogenicity and pathogenicity of SARS-CoV-2 Delta P681R mutation. Nature 602(7896), 300–306 (2022)

[55] Cao, Y., Yisimayi, A., Jian, F., Song, W., Xiao, T., Wang, L., Du, S., Wang, J., Li, Q., Chen, X., Yu, Y., Wang, P., Zhang, Z., Liu, P., An, R., Hao, X., Wang, Y., Wang, J., Feng, R., Sun, H., Zhao, L., Zhang, W., Zhao, D., Zheng, J., Yu, L., Li, C., Zhang, N., Wang, R., Niu, X., Yang, S., Song, X., Chai, Y., Hu, Y., Shi, Y., Zheng, L., Li, Z., Gu, Q., Shao, F., Huang, W., Jin, R., Shen, Z., Wang, Y., Wang, X., Xiao, J., Xie, X.S.: BA.2.12.1, BA.4 and BA.5 escape antibodies elicited by Omicron infection. Nature 608(7923), 593–602 (2022). https://doi.org/10.1038/s41586-022-04980-y

[56] Zhang, X., Wu, S., Wu, B., Yang, Q., Chen, A., Li, Y., Zhang, Y., Pan, T., Zhang, H., He, X.: SARS-CoV-2 Omicron strain exhibits potent capabilities for immune evasion and viral entrance. Signal transduction and targeted therapy 6(1), 430 (2021)

[57] Boulant, S., Stanifer, M., Lozach, P.Y.: Dynamics of virus-receptor interactions in virus binding, signaling, and endocytosis. Viruses 7(6), 2794–2815 (2015). https://doi.org/10.3390/v7062747

[58] Vahey, M.D., Fletcher, D.A.: Influenza a virus surface proteins are organized to help penetrate host mucus. Elife 8, 43764 (2019). https://doi.org/10.7554/eLife.43764

[59] Korosec, C.S., Jindal, L., Schneider, M., Calderon, I., Barca, D., Zuckermann, M.J., Forde, N.R., Emberly, E.: Substrate stiffness tunes the dynamics of polyvalent rolling motors. Soft Matter 17(6), 1468–1479 (2021). https://doi.org/10.1039/D0SM01811B

[60] Wu, C.T., Lidsky, P.V., Xiao, Y., Cheng, R., Lee, I.T., Nakayama, T., Jiang, S., He, W., Demeter, J., Knight, M.G., Turn, R.E., Rojas-Hernandez, L.S., Ye, C., Chiem, K., Shon, J., Martinez-Sobrido, L., Bertozzi, C.R., Nolan, G.P., Nayak, J.V., Milla, C., Andino, R., Jackson, P.K.: SARS-CoV-2 replication in airway epithelia requires motile cilia and microvillar reprogramming. Cell 186(1), 112–130 (2023). https://doi.org/10.1016/j.cell.2022.11.030

[61] Thomson, E.C., Rosen, L.E., Shepherd, J.G., Spreafico, R., da Silva Filipe, A., Wojcechowskyj, J.A., Davis, C., Piccoli, L., Pascall, D.J., Dillen, J.: Circulating SARS-CoV-2 spike N439K variants maintain fitness while evading antibody-mediated immunity. Cell 184(5), 1171–1187 (2021)

[62] Lozach, P.-Y.: Cell Biology of Viral Infections. (2020). https://doi.org/10.3390/cells9112431

[63] Vigerust, D.J., Shepherd, V.L.: Virus glycosylation: role in virulence and immune interactions. Trends in microbiology 15(5), 211–218 (2007). https://doi.org/10.1016/j.tim.2007.03.003

[64] Thaker, S.K., Ch’ng, J., Christofk, H.R.: Viral hijacking of cellular metabolism. BMC biology 17(1), 59 (2019). https://doi.org/10.1186/s12915-019-0678-9

[65] Marques-Pereira, C., Pires, M.N., Gouveia, R.P., Pereira, N.N., Caniceiro, A.B., Rosário-Ferreira, N., Moreira, I.S.: SARS-CoV-2 Membrane Protein: From Genomic Data to Structural New Insights. International Journal of Molecular Sciences 23(6) (2022). https://doi.org/10.3390/ijms23062986

[66] Shen, L., Bard, J.D., Triche, T.J., Judkins, A.R., Biegel, J.A., Gai, X.: Emerging variants of concern in SARS-CoV-2 membrane protein: a highly conserved target with potential pathological and therapeutic implications. Emerging Microbes and Infections 10(1), 885–893 (2021). https://doi.org/10.1080/22221751.2021.1922097

[67] Pearson, J., Wessler, T., Chen, A., Boucher, R.C., Freeman, R., Lai, S.K., Pickles, R., Forest, M.G.: Modeling identifies variability in SARS-CoV-2 uptake and eclipse phase by infected cells as principal drivers of extreme variability in nasal viral load in the 48 h post infection. Journal of theoretical biology 565(March), 111470 (2023). https://doi.org/10.1016/j.jtbi.2023.111470

[68] Callaway, E.: Beyond Omicron: What’s next for SARS-CoV-2 Evolution. Nature 600, 204–207 (2021)

[69] Harari, S., Tahor, M., Rutsinsky, N., Meijer, S., Miller, D., Henig, O., Halutz, O., Levytskyi, K., Ben-Ami, R., Adler, A., Paran, Y., Stern, A.: Drivers of adaptive evolution during chronic SARS-CoV-2 infections. Nature Medicine 28(7), 1501–1508 (2022). https://doi.org/10.1038/s41591-022-01882-4

[70] Chakraborty, C., Sharma, A.R., Bhattacharya, M., Lee, S.-S.: A detailed overview of immune escape, antibody escape, partial vaccine escape of SARS-CoV-2 and their emerging variants with escape mutations. Frontiers in Immunology 13, 801522 (2022)

[71] Ao, D., Lan, T., He, X., Liu, J., Chen, L., Baptista-Hon, D.T., Zhang, K., Wei, X.: SARS-CoV-2 Omicron variant: Immune escape and vaccine development. MedComm 3(1), 126 (2022)

[72] Thorne, L.G., Bouhaddou, M., Reuschl, A.K., Zuliani-Alvarez, L., Polacco, B., Pelin, A., Batra, J., Whelan, M.V.X., Hosmillo, M., Fossati, A., Ragazzini, R., Jungreis, I., Ummadi, M., Rojc, A., Turner, J., Bischof, M.L., Obernier, K., Braberg, H., Soucheray, M., Richards, A., Chen, K.H., Harjai, B., Memon, D., Hiatt, J., Rosales, R., McGovern, B.L., Jahun, A., Fabius, J.M., White, K., Goodfellow, I.G., Takeuchi, Y., Bonfanti, P., Shokat, K., Jura, N., Verba, K., Noursadeghi, M., Beltrao, P., Kellis, M., Swaney, D.L., García-Sastre, A., Jolly, C., Towers, G.J., Krogan, N.J.: Evolution of enhanced innate immune evasion by SARS-CoV-2. Nature 602(7897), 487–495 (2022). https://doi.org/10.1038/s41586-021-04352-y

[73] Jian, F., Yu, Y., Song, W., Yisimayi, A., Yu, L., Gao, Y., Zhang, N., Wang, Y., Shao, F., Hao, X., Xu, Y., Jin, R., Wang, Y., Xie, X.S., Cao, Y.: Further humoral immunity evasion of emerging SARS-CoV-2 BA.4 and BA.5 subvariants. (2022). https://doi.org/10.1016/S1473-3099(22)00642-9

[74] Baccam, P., Beauchemin, C., Macken, C.A., Hayden, F.G., Perelson, A.S.: Kinetics of Influenza A Virus Infection in Humans. Journal of Virology 80(15), 7590–7599 (2006). https://doi.org/10.1128/jvi.01623-05

[75] Agostini, M.L., Andres, E.L., Sims, A.C., Graham, R.L., Sheahan, T.P., Lu, X., Smith, E.C., Case, J.B., Feng, J.Y., Jordan, R., Ray, A.S., Cihlar, T., Siegel, D., Mackman, R.L., Clarke, M.O., Baric, R.S., Denison, M.R.: Coronavirus susceptibility to the antiviral remdesivir (GS-5734) is mediated by the viral polymerase and the proofreading exoribonuclease. mBio 9(2) (2018). https://doi.org/10.1128/mBio.00221-18

[76] Ghosh, S., Dellibovi-Ragheb, T.A., Kerviel, A., Pak, E., Qiu, Q., Fisher, M., Takvorian, P.M., Bleck, C., Hsu, V.W., Fehr, A.R., Perlman, S., Achar, S.R., Straus, M.R., Whittaker, G.R., de Haan, C.A.M., Kehrl, J., Altan-Bonnet, G., Altan-Bonnet, N.: β-Coronaviruses Use Lysosomes for Egress Instead of the Biosynthetic Secretory Pathway. Cell 183(6), 1520–1535 (2020). https://doi.org/10.1016/j.cell.2020.10.039

[77] Heffernan, J.M., Smith, R.J., Wahl, L.M.: Perspectives on the basic reproductive ratio. Journal of the Royal Society Interface 2(4), 281–293 (2005). https://doi.org/10.1098/rsif.2005.0042

